# Style deflection is determined by the handedness of phyllotaxis and auxin-induced differential cell elongation in a species with mirror-image flowers

**DOI:** 10.1101/2024.06.14.598852

**Authors:** Caroline Robertson, Haoran Xue, Marco Saltini, Alice L. M. Fairnie, Dirk Lang, Merijn H. L. Kerstens, Viola Willemsen, Robert A. Ingle, Spencer C. H. Barrett, Eva E. Deinum, Nicola Illing, Michael Lenhard

## Abstract

Many animals and plants show left-right (LR) asymmetry. In some animal systems, handedness has a simple genetic basis, which has allowed identifying how handedness is determined at the molecular level, even if its functional relevance often remains unclear. Mirror-image flowers represent an example of LR asymmetry of clear functional significance, with the reciprocal placement of male and female organs in left- versus right-handed flowers promoting cross-pollination. Here, we use the South African geophyte *Cyanella alba* to study how handedness of its mirror-image flowers is determined and elaborated during development. Inflorescences of *C. alba* produce flowers with a largely consistent handedness. However, we find that this handedness has no simple genetic basis, and individual plants can switch their predominant handedness between years. Rather, it is the direction of the phyllotactic spiral that determines floral handedness. Cellular analysis combined with biophysical modelling demonstrates that style deflection is driven by increased cell expansion in the adaxial carpel facing the next oldest flower compared to the other adaxial carpel. The more expanding carpel shows transcriptional signatures of increased auxin signaling compared to the less expanding one, and auxin application to the latter can reverse the orientation of style deflection. We propose that a recently described inherent LR auxin asymmetry in the initiating organs of spiral phyllotaxis determines handedness in *C. alba*, representing a conserved non-genetic mechanism for creating a stable floral polymorphism. This mechanism links chirality across different levels of plant development and exploits a developmental constraint in a core patterning process to produce morphological variation of ecological relevance.

## INTRODUCTION

Left-right (LR) asymmetry is a fascinating feature of many plants and animals. Striking examples include the asymmetric placement of internal organs in vertebrates and the left versus right coiling of snail shells ^1,2^. Such LR asymmetries raise three fundamental questions. First, how is symmetry broken in a consistent manner to determine left and right? Second, how is this translated, from genes to cellular and tissue level processes, to produce asymmetric morphologies? Third, what is the biological function of LR asymmetry? The first two questions are being answered for the LR asymmetry of internal organs in vertebrates and shell coiling direction in snails ^3–8^. In both systems the inherent chirality of different cytoskeletal elements appears to be exploited for the initial symmetry breaking event, which is then translated into an asymmetric morphology via a partially conserved pathway. In mouse embryos, inherently chiral motile cilia on the cells in the node cause a left-ward flow of extracellular fluid that breaks symmetry ^5,6,9^. In *Lymnaea* snails a genetic polymorphism determines coiling direction, with the dominant *D* allele, which encodes a nucleator for actin filaments, causing right-handed coiling ^3,7,10^. In both cases LR-asymmetric upregulation of *Nodal* and *Pitx2* expression translates the initial symmetry breaking event into an asymmetric morphology. Despite this progress in understanding the molecular basis of handedness, the functional relevance of LR asymmetry in these examples remains largely unclear.

Several examples of LR asymmetry and left- or right-handed helical growth also exist in plants^2^. These include the left-right asymmetry of leaves in several species ^11^, left- or right-handed helical arrangement of leaves and flowers around the stem (i.e. phyllotaxis) ^12^, the twining of tendrils ^13^, and the left- or right-handed helical growth of mutants ^14^. Several important components that underlie helical growth *per se* have been identified, such as a polar auxin transport-based mechanism that determines the initiation of organs at the shoot meristem in a phyllotactic spiral and indirectly seems to cause LR asymmetries in leaves ^15,16^, or gelatinous G-fibres in twisting tendrils ^17^. But how the handedness of the resulting helices is determined remains unknown. The important exception are mutants with consistently left- or right-handed helical growth ^14^. Most of these result from mutations in tubulin or microtubule-associated proteins that cause a spiral arrangement of cortical microtubules, with the handedness of this spiral determining the handedness of helical growth at the cell or organ level ^18,19^. These mutants underscore the value of genetic polymorphisms causing helical growth with a consistent orientation for identifying the molecular determinants of handedness.

Mirror-image flowers (enantiostyly) represent a case of LR asymmetry in plants for which a clear functional relevance has been demonstrated and where - at least in some taxa - the handedness of the flowers is genetically determined ^20–22^. This provides an opportunity to identify both the molecular and developmental control of handedness and its adaptive significance ^23^. In enantiostylous flowers the style is deflected to the left- or right-side of the midline that runs along the dorsoventral axis of the flower ^24^, and in most species a pollinating anther is deflected in the opposite direction (Fig. 1). Enantiostylous species are classified based on the distribution of left- and right-styled flowers on individuals ^24^. In monomorphic enantiostyly, both forms of flower can be found on a single individual, whereas in dimorphic enantiostyly a plant only produces left- or right-styled flowers, called L-morph or R-morph plants, respectively.

**Figure 1:**
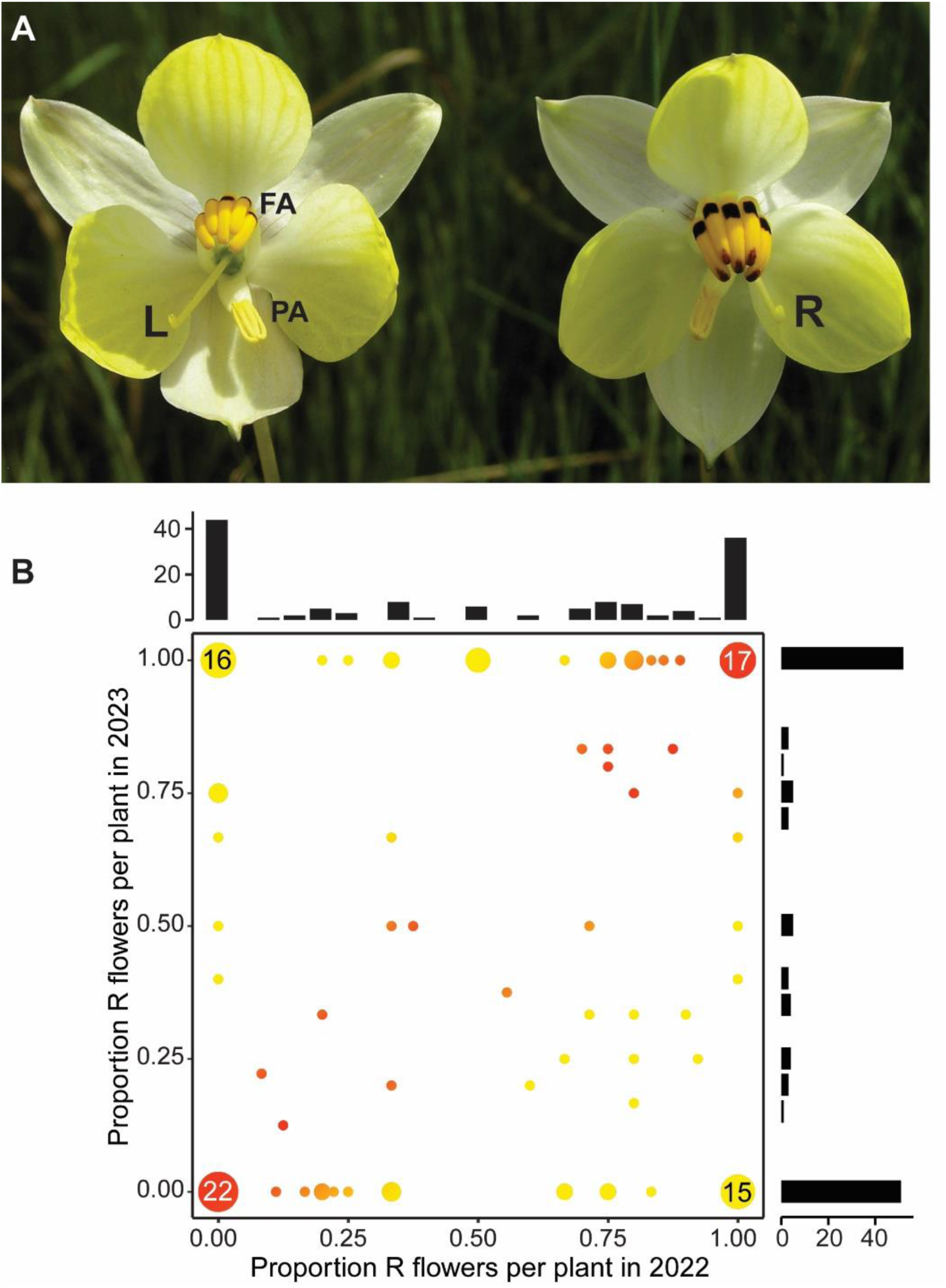
Floral handedness has no simple genetic basis in *C. alba*. (**A**) Image of L- and R-flowers of *Cyanella alba* subsp*. flavescens.* Feeding anthers (FA) and pollinating anther (PA) are indicated. L and R indicate left- and right-deflected styles. (**B**) Distribution of floral phenotypes of the same cohort of 135 plants in 2022 and 2023. The colour gradient indicates the consistency in floral orientation between years (red: fully consistent, yellow: fully inconsistent). Black bars on the outside show marginal distributions.

Enantiostyly is reliably reported from 11 unrelated angiosperm families and represents a striking case of convergent evolution in form and function. Most enantiostylous species are of the monomorphic type ^23,24^, with dimorphic enantiostyly reliably reported from just two monocotyledonous lineages: Pontederiaceae (*Heteranthera*) and Haemodoraceae (*Wachendorfia*, *Barberetta*). A third instance in Tecophilaeaceae (*Cyanella*) has been reported ^20^, but this has not been fully confirmed. Experimental studies have demonstrated that enantiostyly represents a floral adaptation that promotes cross-pollination and reduces sexual interference between male and female function in hermaphroditic flowers ^25,26^. It does so by optimizing the placement of pollen on the bodies and wings of pollinating insects for efficient disassortative pollen transfer to styles on the opposite morph. Studies using genetic markers indicate that dimorphic enantiostyly promotes significantly higher levels of outcrossing than the monomorphic condition ^21,27^. The efficiency of dimorphic enantiostyly in promoting disassortative pollination has also been demonstrated using fluorescently labelled pollen grains in *Wachendorfia paniculata* ^28^. Similar studies in the sister genus, *Barberetta aurea* reported that transfer of fluorescent dye occurred mainly between reciprocally positioned floral organs ^29^. Thus, at least for some species/pollinator combinations the pollen that left-styled (L) flowers deposit on the right side of visiting insects is transferred efficiently to the stigmas of right-styled (R) flowers, and vice versa. The mode of inheritance of dimorphic enantiostyly has only been determined in *Heteranthera multiflora* (now *H. missouriensis*), where handedness is determined by a genetic polymorphism segregating in a Mendelian manner ^22^. Plants that carry the dominant allele at the polymorphic locus form R flowers, while the homozygous recessive genotype produces L flowers.

Here, we investigate the molecular and development basis of enantiostyly in *Cyanella alba subsp. flavescens* (hereafter *C. alba*), a long-lived, deciduous geophyte restricted to the Western and Northern Cape regions of South Africa ^30^. Plants form deep-seated corms that produce a basal rosette of narrow erect leaves after winter rains with flowering occurring from early August to early October. The inflorescence is a compact raceme with very little stem elongation, and flowers are borne individually on long pedicels (10-20 cm), giving the appearance of solitary flowers emerging from the rosette. Each plant produces a small number (generally not more than 15) of long-lived flowers during the flowering season, but daily display size is most commonly one to two flowers ^31^. *C. alba* is a candidate for showing dimorphic enantiostyly based on previous work ^20,24,32^, as the majority of plants were reported to form flowers of only one handedness and only a few plants produced both types of flowers ^32^. Thus, floral handedness in *C. alba* may have a genetic basis.

Our investigation used a sequencing-based approach to search for the presumed genetic basis of floral handedness in *C. alba*. By analogy to other similar floral polymorphisms and to enantiostyly in *H. missouriensis*, we hypothesized that floral handedness in *C. alba* was under the control of a single *ENANTIOSTYLY* (*E*) locus with a dominant and recessive allele. Under such a scenario, disassortative (intermorph) mating between L- and R-morphs would produce an equal ratio of L- and R-morphs in the progeny, with one morph homozygous recessive and the other heterozygous at the *E* locus. Our study aimed to answer three questions. First, is there indeed a genetic polymorphism that determines floral handedness in *C. alba*? Second, what is the symmetry-breaking event that determines the direction of style deflection in flowers? Third, which cellular and developmental processes cause consistent style deflection to one side?

## RESULTS

### Sequencing-based analysis and transcriptomics do not support a simple genetic polymorphism for floral handedness

To determine whether there is a genetic polymorphism determining handedness, we established a reference genome assembly from an individual that had formed seven consistently R-flowers in 2021 (see Supporting Information Text). We performed Illumina whole-genome sequencing on 25 *C. alba* individuals that had four or more flowers of consistent handedness in 2021 (15 L and 10 R), and 20 with six or more flowers of consistent handedness in 2022 (11 L and 9 R; see Supporting Information Text). In addition, we performed RNA-sequencing of dissected styles from bud stages covering the critical time window when the orientation of style deflection is determined (see Supporting Information Text). Using this data, we asked whether (i) there is a bi-allelic locus associated with style orientation; (ii) there is a hemizygous region determining handedness, as demonstrated in several heterostylous species ^33–35^; or (iii) whether there are any transcripts expressed exclusively in plants with one style orientation. To answer these questions, we performed (i) a genome-wide association study (Table 1), (ii) searched for a genomic region exclusively present in plants of one style orientation using both coverage- and *k*-mer based approaches (Figure S1, Table S1), and (iii) identified differentially expressed genes between plants with the different style orientations. However, as detailed in the Supporting Information Text, none of these approaches provided any convincing evidence for a simple genetic polymorphism underlying floral handedness in *C. alba*. For example, no orientation-associated single-nucleotide polymorphisms (SNPs) were shared between the datasets from the two years and the combined dataset (Table 1), and the *k*-mer based approach identified more left-associated *k*-mers in the 2021 dataset, but many more right-associated ones in the 2022 dataset (Figure S1).

**Table 1:** Numbers of morph-associated SNPs in *Cyanella alba* subsp*. Flavescens*. The number of morph-associated SNPs in each dataset where 80% or more individuals of a given handedness are heterozygous at the SNP site is given. The analysis was restricted to the protein-coding regions of the *C. alba flavescens* genome.

|  | SNPs where L-plants are heterozygous | SNPs where R-plants are heterozygous |
| --- | --- | --- |
| Illumina2021 dataset (25 individuals) | 10 | 12 |
| Illumina2022 dataset (20 individuals) | 12 | 17 |
| Combined dataset (45 individuals) | 0 | 1 |

### Switching floral handedness on individual plants between flowering seasons indicates that the direction of style deflection is not genetically determined

The above conclusion that there is unlikely to be a genetic basis to handedness is further supported from our demographic studies of individually marked plants during the flowering seasons of 2022 and 2023. In 2022, 188 plants in three populations were permanently marked with metal pins, and in 2023 we recovered 135 of these plants. We followed the plants throughout the duration of both flowering seasons and recorded the number of L- and R- flowers produced by each individual (Figure 1D). Strikingly, of the 44 plants recovered in 2023 that had formed exclusively L-flowers in 2022, 22 retained their handedness and 22 switched handedness in 2023. Of the latter, 16 produced exclusively R-flowers and 6 produced both L- and R- flowers (proportion of R flowers within a plant ranged from 40-75%). Conversely, of the 36 plants recovered in 2023 that had exclusively R-flowers in 2022, 19 switched their handedness in 2023. Again, the same trend was seen as above with the majority, 15 plants, producing exclusively L-flowers in 2023, and the remaining four plants (21%) producing both flower types (proportion of L flowers within a plant ranged from 33-60%). Thus, these results demonstrate that the handedness of flowers produced by an individual plant during one flowering season is highly consistent, but that this consistent handedness can switch within an individual from one year to the next. This supports the above conclusion that floral handedness in *C. alba* is not determined by a genetic polymorphism and suggests that the species exhibits monomorphic enantiostyly and not dimorphic enantiostyly as earlier assumed.

### The handedness of the phyllotactic spiral determines the consistent bias in floral handedness within plants

The lack of direct genetic control of handedness raises an intriguing question: How can flowers in an inflorescence consistently develop an L or R morphology without genetic input, and how can they switch their annually consistent bias for one side to the other in the next year? One developmental process with a highly consistent handedness over the growing period of an inflorescence is the direction of the phyllotactic spiral. Its handedness is also known to cause LR asymmetries in leaves ^15,16^. *C. alba* shows spiral phyllotaxis (Figure S2), with organs initiated at the shoot apical meristem in either a clockwise or counter-clockwise direction. Changes in the handedness of the phyllotactic spiral produced by one shoot meristem are exceedingly rare ^36^, providing stability in any one year. This also holds true in *C. alba*, as none of 133 plants we sampled showed a consistent handedness reversal of the phyllotactic spiral. Moreover, studies of many angiosperm species indicate that the handedness of the phyllotactic spiral is not under genetic control, and that initiating shoot meristems randomly establish either a clockwise or counter-clockwise spiral with equal probability ^12^. This pattern is also true for the individual lateral meristems on a single plant. Thus, it is likely that the shoot meristems initiated in successive years by *C. alba* randomly adopts a clockwise or counter-clockwise orientation. We therefore asked whether the direction of style deflection in *C. alba* flowers is predicted by the handedness of the phyllotactic spiral.

To address this question, we determined the handedness of the phyllotactic spiral on 132 *C. alba* plants in the 2023 flowering season and the phenotype of between 1 and 7 open flowers and sufficiently mature buds for recognizing the style orientation per plant (average 2.74, median 3 flowers per plant; a small number of flowers with unclear style orientation were excluded). This revealed a strong correlation between the direction of the phyllotactic spiral and floral handedness, with a correlation coefficient of 0.82 (0.76 – 0.87). Most plants with a counter-clockwise or right-handed phyllotactic spiral (defined from older to younger organs) formed only R flowers, and *vice versa* for plants with a clockwise or left-handed phyllotactic spiral. Twenty plants produced both R and L flowers, and there were very few plants (5/132) where floral handedness consistently contradicted that predicted by the phyllotaxis direction. Very similar results were observed in 53 plants analyzed in 2024 (Figure S2J). Thus, the handedness of the phyllotactic spiral in *C. alba* determines floral handedness, suggesting that it provides the consistently biased input into flower development to ensure the non-random distribution of style deflection of individual plants through time.

We next asked whether deviations from the floral handedness predicted by phyllotaxis were randomly distributed in an inflorescence. For this we plotted the fraction of such deviations per absolute flower position in an inflorescence and compared it to the overall average of all flowers. The distribution of deviations was non-random across the different positions (*p*<0.0001, *Χ*^2^-test), with higher deviation rates at the very first and at higher positions beyond position 5, and the lowest deviation rates at positions 2 to 4, i.e., the very first and the later flowers of the season were more likely to have deviating handedness. The reason for this non-random distribution of errors is unclear at present, but it may suggest an additional flower-flower interaction that cannot operate on the first flower and becomes weaker over time.

### Style deflection results from differential elongation of the two adaxial carpels

The styles of *C. alba* are initially straight with no discernible difference between L- and R- flowers (Figure 3A). When styles of early buds reach 70% or more of the length of the pollinating anther, they begin to deflect slightly to the left or right of the midline. The stigma tip can also be slanted at this stage, with the slant facing the direction to which the style will be deflected. In mid-stage buds, one to two days before anthesis, the style is clearly deflected from the midline, and in open flowers the style and pollinating anther are fully deflected to opposite sides of the flower. The style is relatively straight, with the exception of the stigma tip curving sharply upwards. The style movement to one side can be observed using time-lapse microscopy in dissected flowers cultivated *in vitro* (Video S1). The hinge-like motion of the style is coupled with expansion and elongation of the ovary, as confirmed by measuring the distances between the three marked spots on the ovary, which increased by 14% and 10% in the length and width direction over the course of the video, respectively (Video S1). These observations suggested that style deflection is driven by differential elongation of the carpels. The *C. alba* gynoecium consists of three carpels, with the two adaxial carpels lying on either side of the dorsoventral midline of the flower and the third carpel below, such that the midline bisects it (Figure 3B). If one of the two adaxial carpels elongated more than the other, this could push the style away from the more strongly elongating carpel.

**Figure 2:**
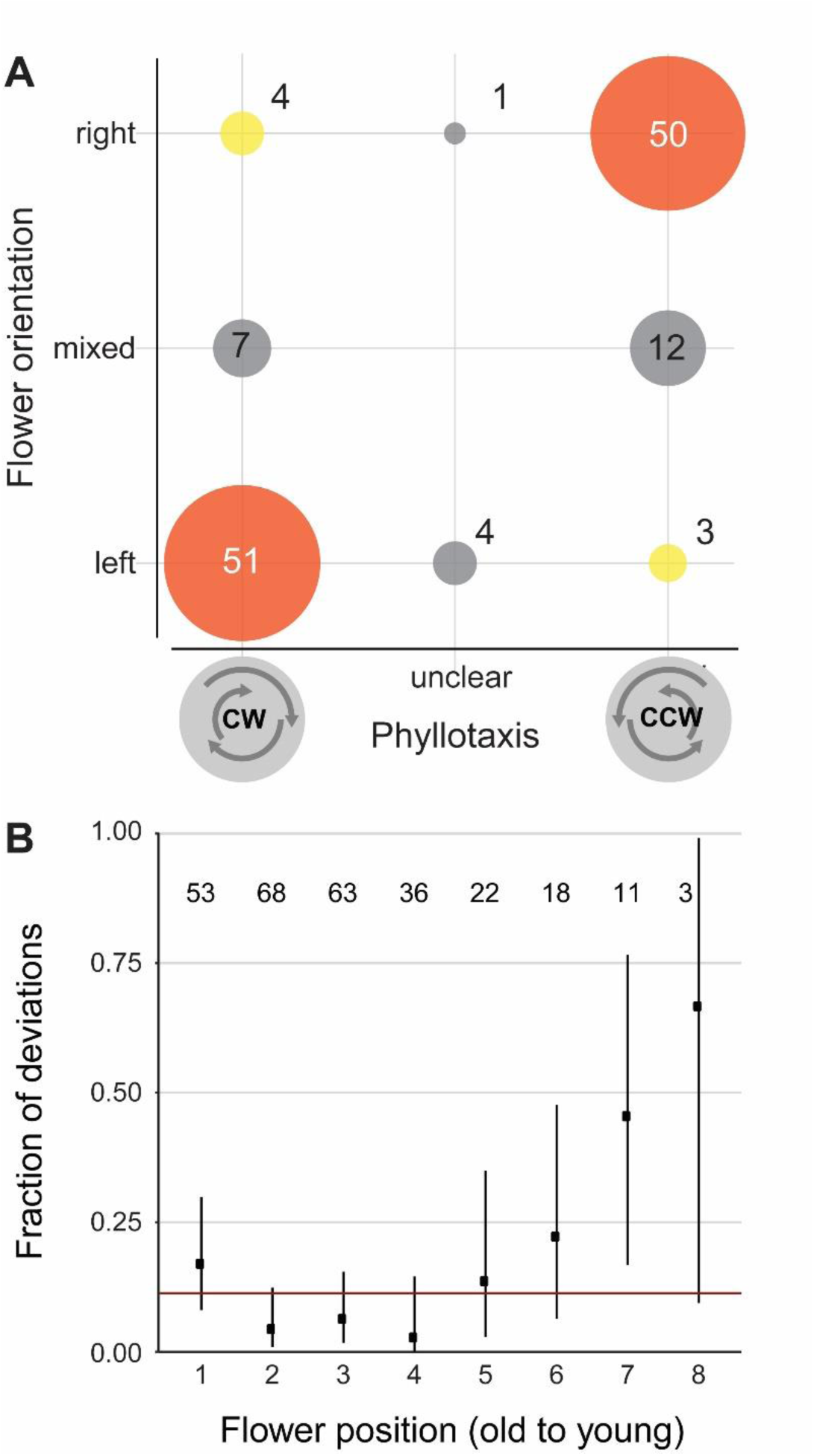
Direction of the phyllotactic spiral determines floral handedness in *Cyanella alba* subsp*. Flavescens*. (**A**) Correlation of phyllotactic direction (CW: clockwise, CCW: counter-clockwise) and floral handedness. Numbers in the diagram indicate the number of individual plants. ‘Unclear’ phyllotaxis indicates plants with individual divergence angles that are inconsistent with their overall phyllotaxis orientation. See Materials and Methods in Supporting Information for more details and our interpretation of these cases. (**B**) Distribution of deviations from main pattern (flowers with the opposite style deflection compared to the orientation predicted based on phyllotaxis) across flower positions in an inflorescence. Numbers in diagram indicate the number of flowers scored at each position. Horizontal red line shows average fraction of such errors over all flower positions. Dots are observed values and vertical lines indicate 95% confidence intervals for the fraction of deviations per position calculated based on a binomial test.

**Figure 3:**
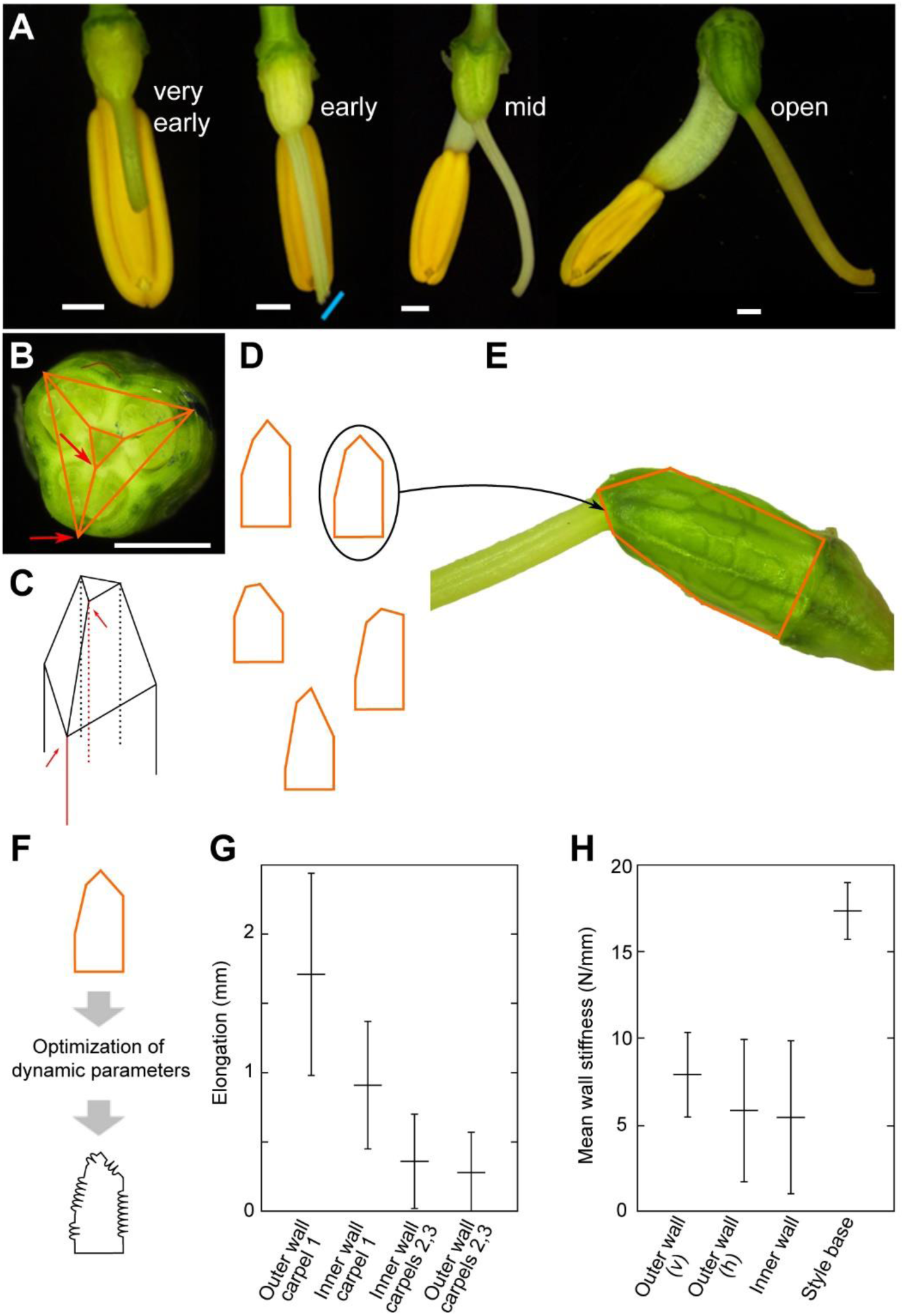
Schematic of the data-driven and mechanistic models and analysis of ovaries of *Cyanella alba* subsp*. flavescens*. (**A**) Stereomicroscope images of *C. alba* buds at four stages of development reveal the developmental sequence of stylar bending. Blue line indicates the slant of the stigma in early-stage buds. Scale bars are 1 mm. (**B**) Cross-section of a *C. alba* ovary indicating the trilocular architecture. Orange lines represent a cross-section of the modelling polyhedron. Red arrows indicate the edges of the polyhedron representing inner and outer wall of a carpel. The outer-wall positions overlie the carpel midveins. (**C**) The ovaries are modelled as polyhedra composed of triangular prisms topped with a truncated pyramid. Red edges denote the inner and outer walls of the same carpel highlighted in (**B**). (**D**) Two-dimensional projection of the lateral view of a collection of polyhedra. (**E**) The selection process of the best fitting polyhedron for modelling the ovaries of *C. alba*, consists of choosing among all possible polyhedra based on area, perimeter, and length and width of the ovary. (**F**) Biophysical parameters of our bead-spring models are optimised to achieve an accurate fit of the polyhedron to the ovary structure. (**G**, **H**) Elongation of the inner and outer walls of the C. alba ovaries (**G**) and stiffness of the organ walls (**H**) as predicted by our mechanistic model. Stiffness is shown in the vertical (v) and horizontal (h) directions. Error bars represent standard deviation. n=6

To test the feasibility of this mechanism and investigate the contribution of differential elongation of different carpel walls, we developed a biophysical model of the *C. alba* pistil. Our aim was to test whether differential elongation of different carpels can drive style deflection to the extent observed in *C. alba* ovaries. As individual ovaries varied substantially in size and shape, we opted for a model that captures organ shape in a simple description, to which we could fit the individual ovaries, rather than a model reproducing realistic organ shapes. To this end, we developed a bead-spring model with as few springs per carpel as possible to capture the essence of the developmental process ^37^. Bead-spring mechanical models are well-suited for identifying differential growth patterns without the need for complex geometries or full organ-level modelling ^38^. We used a three-dimensional polyhedral model to represent the ovary, which is composed of a triangular prism topped by a truncated pyramid (Figure 3C and Supporting Information Text; Figures S3, S4; Tables S3 to S8). Deflection of the truncated face of the pyramid such that it is no longer parallel to the base of the triangular prism indicates style deflection. We applied this model to images of six ovaries in two steps. First, we identified for each ovary the “target” polyhedron whose two-dimensional side view most accurately represented the ovary in its later stages of development (see Figure 3B-E, and Figures S3, S4). The resulting solid figures were skewed at the top, reflecting the differential carpel elongation responsible for stylar deflection. Second, we searched for the parameters that most closely reproduced the target polyhedron as a bead-spring system at its dynamic equilibrium, with the polyhedron edges as springs (Figure 3F). We allowed different elongation (increase in resting length) and stiffness in the springs on the carpel 1 (Figure 4A; the adaxial carpel opposite the direction of style deflection) vs carpel 2 (the adaxial carpel on the side to which the style deflects) and carpel 3 (the abaxial carpel) sides compared to a symmetric polyhedron based on very-early bud measurements (Figure S5A,B, Table S5 and Supporting Text). We used random search optimisation algorithms ^39^, to identify the parameters of our mechanistic model that best reproduced the target polyhedron and, hence, of the ovary of *C. alba* during the late developmental stage (Figures 3, S5, S6, Table S7).

**Figure 4:**
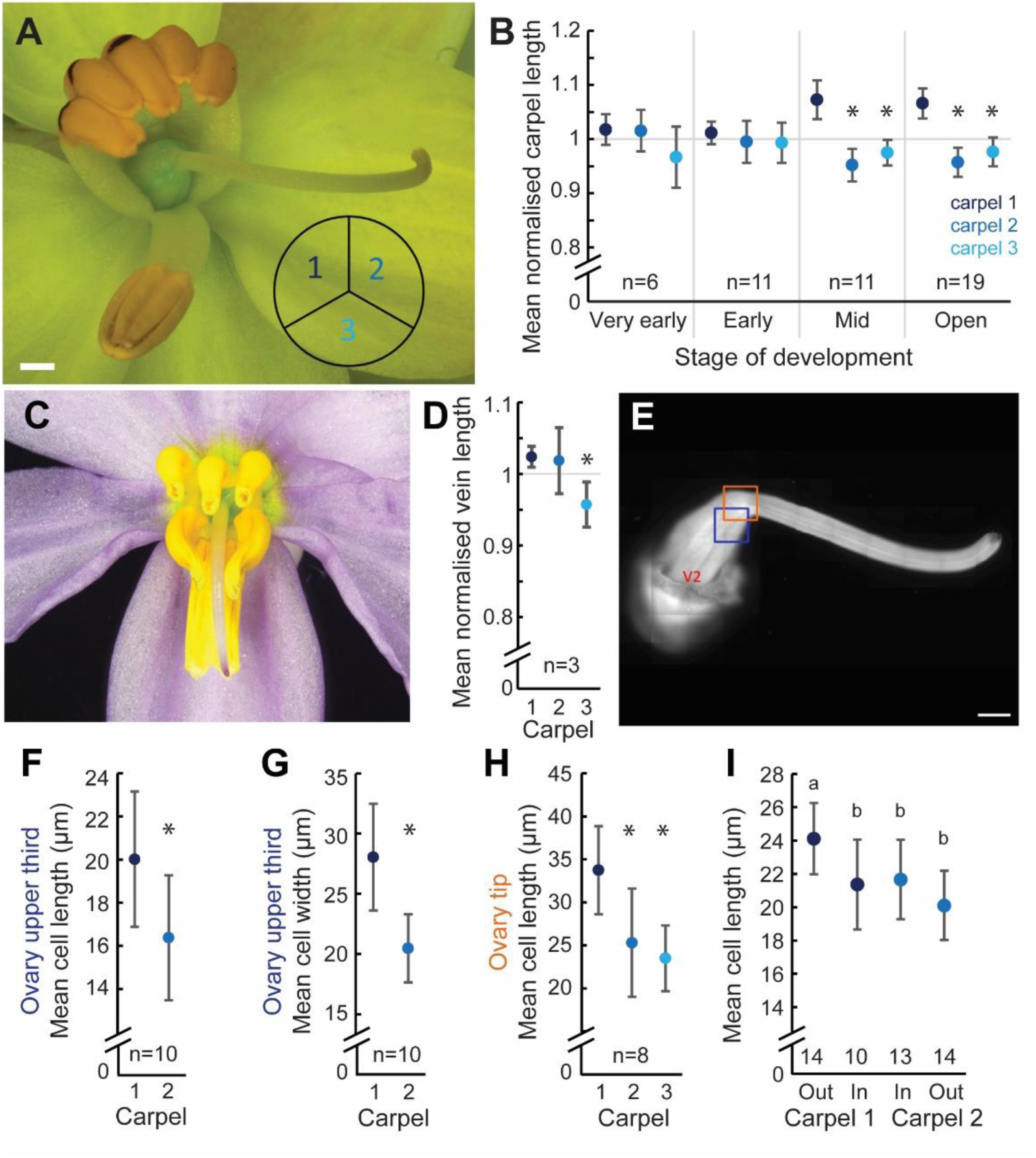
Differential carpel elongation underlies style deflection in *Cyanella alba* subsp*. Flavescens*. (**A**) An open *C. alba* flower. The arrangement of the three carpels in the trilocular ovary is indicated by the subdivided circle. (**B**) Measurements of carpel lengths from ovaries of flowers at the indicated stages of development. Lengths were normalised to the average value of the three carpels per ovary. Values are means ± SD here and throughout the other panels. Sample sizes are indicated and show the number of ovaries from which the paired measurements were taken. *: significantly different from carpel 1 (p<0.05, paired t- test). (**C**) *C. hyacinthoides* is a closely related speci*es* without enantiostyly. The style is deflected downwards and placed centrally between the six anthers. (**D**) Measurements of carpel lengths from *C. hyacinthoides* ovaries. *: significantly different from carpel 1 (*p*<0.05, paired *t*-test). (**E**) Fluorescence micrograph of a calcofluor white-stained pistil. Blue and orange boxes indicate the regions where the measurements of cell sizes shown in (**F**, **G**) and (**H**) were taken, respectively. V2 indicates midvein of carpel 2. (**F**, **G**) Cell lengths (**F**) and cell widths (**G**) of epidermal cells overlying the midveins of carpels 1 and 2 in the upper third of the ovary are shown. Cell length is the extension of the cell along the long axis of the ovary, and cell width the extension perpendicular to the long axis of the ovary. *: significantly different from carpel 1 (p<0.0001, paired t-test). (**H**) Cell lengths of epidermal cells overlying the midveins of carpels 1 and 2 at the tip of the ovary are shown. *: significantly different from carpel 1 (p<0.0001, paired t-test). (**I**) Lengths of subepidermal cells in the outer and inner walls of carpels 1 and 2. The number of carpels from which the measurements were taken is indicated inside the graph. Letters indicate mean cell lengths that are significantly different as determined by a two-way repeated measures ANOVA followed by Tukey’s HSD post-hoc test (α=0.05). In total 14 ovaries were dissected and imaged, but for some of them individual walls could not be imaged in sufficient quality to allow cell-length measurements. Scale bars are 1 mm.

The results of this optimization indicated that both the inner and outer walls of carpel 1 undergo more significant elongation than those of carpels 2 and 3 (Figure 3G). The slight elongation seen in carpels 2 and 3, as predicted by our model, arises from the tensile forces generated by links between different carpels at the base of the style. This effect is driven by the greater relative stiffness of the style base compared to other system walls (Figure 3H) and contributes to guiding the shape and morphogenesis of the style ^40,41^. Our analysis revealed that both inner and outer walls of carpel 1 end up being compressed (actual length shorter than spring resting length), whereas those in carpels 2 and 3 are stretched (Table S8). Based on the analyses of six *C. alba* ovaries through our models, the primary driver behind the elongation of carpel 1 is the extension of the outer wall, with an average detected elongation relative to the initial length of the early bud stage of 2.47 (SD = 0.41) times, as opposed to the inner wall mean relative elongation of 1.28 (SD = 0.18) times (Figure 3G).

The above model makes two key predictions. First, the outer wall of carpel 1 should elongate more than the outer wall of carpel 2. Thus, in an R-flower we would predict the top left carpel (when viewed from the front) would elongate more than the top right carpel (Figure 4A). The second prediction is that the outside wall of carpel 1 should elongate more than the inner wall corresponding to the placenta. To test the first prediction, we measured the lengths of the three carpels in the ovary along their prominently visible midveins at four different stages of development and combined data for L- and R-flowers. Carpel 1 was significantly longer than carpel 2 and the abaxial carpel 3 in both open flowers and mid-stage buds (Figure 4B). Carpel 2 was on average 89.9% ± 4.3% and 88.9% ± 5.42% of the length of carpel 1 in open flowers and mid-stage buds, respectively. By contrast, carpel lengths were indistinguishable in early or very early-stage buds (Figure 4B). This mirrors the observation that clear deflection of the style to the left or right is only visible in mid-stage buds and open flowers and supports the hypothesis that differential elongation between the two adaxial carpels causes style deflection in *C. alba*.

To further support the link between differential carpel elongation and style deflection, we repeated this experiment on *C. hyacinthoides*, a close relative of *C. alba* that does not exhibit enantiostyly (Figure 4C). Styles of *C. hyacinthoides* are not deflected to the left or right, but instead towards the abaxial side. The structure of the trilocular *C. hyacinthoides* ovary matches that of *C. alba*. Carpels 1 and 2 of *C. hyacinthoides* were indistinguishable in length, but carpel 3 was significantly shorter than carpel 1 (Figure 4D). This suggests that the reduced expansion of carpel 3 underlies the downward deflection of the style in *C. hyacinthoides*, whereas the differential elongation of carpels 1 and 2 determines the LR deflection of the style in *C. alba*.

We next asked whether the stronger elongation of carpel 1 in *C. alba* was solely due to more cell elongation or was also accompanied by more cell division. The length of the epidermal cells overlying the carpel midveins was measured from confocal laser-scanning microscopy images of the distal third of ovaries (Figure 4E, Figure S7). Cell length (i.e. the size of the cell along the ovary’s longitudinal axis) was significantly greater along the midvein of carpel 1 than along carpel 2, with mean cell length in carpel 2 only 81.8% of that observed in carpel 1 (Figure 4F). This difference is more than sufficient to explain the observation that carpel 2 length is on average 89.9% of carpel 1 length (Figure 4B). In addition, carpel 1 cells are significantly wider than carpel 2 cells (Figure 4G). To confirm the observed differences in cell length, we repeated the measurement at the tip of the ovary where it transitions into the style. Again carpel 1 cells were significantly longer than carpel 2 (and also carpel 3) cells, with the difference between carpels 1 and 2 even greater (25%) than in the upper third of the ovary (Figure 4H), suggesting a proximo-distal gradient in differential cell elongation.

To test the second prediction from our model, we compared cell lengths between the outer carpel walls and the inner walls at the placenta for carpels 1 and 2 (Figure S8). Subepidermal cells were used to test this prediction, as they could be clearly and reproducibly measured, and the epidermis was often damaged due to the need to remove the ovules before imaging. As expected, outer-wall cells from carpel 1 were longer than outer-wall cells from carpel 2, confirming the above measurements. Outer-wall cells from carpel 1 were also longer than inner-wall cells from carpel 1, while those of carpel 2 were not (Figure 4I). Thus, these measurements fully support the predictions from the model. Collectively our empirical results and biophysical model indicate that stronger isotropic cell expansion in the outer wall of carpel 1 than carpel 2 drives style deflection to the opposite side of the midline from carpel 1.

### Differential carpel growth is associated with increased expression of auxin-induced and elongation-related genes in carpel 1

Given the differential elongation of the two adaxial carpels, we compared their gene expression patterns using RNA-seq data. Outer walls of carpel 1 and 2 were dissected from early- and mid-stage buds with their individual directions of style deflection clearly discernible. We identified 53 differentially expressed transcripts (FDR<0.05, absolute log2 fold-change(carpel 1/carpel2) >1). These included four *Auxin/Indole-3-Acetic Acid* (*Aux/IAA*) and one *SMALL AUXIN UP-REGULATED* (*SAUR*) transcripts, all of which were upregulated in carpel 1 (Table S9). To complement this individual-transcript based analysis, we functionally annotated transcripts using Mercator4 ^42^ and tested for bins that differed significantly from the remaining transcriptome in terms of their distribution of log2 fold-change values. In total, 106 bins showed a significantly different distribution compared to the background (Table S10). The empirical cumulative-distribution function (ECDF) plots indicated that for many of these bins gene-expression differences between the two carpels were less variable than the transcriptomic background, consistent with these being large bins of house-keeping genes (Figure S9). We identified three bins with a possible link to cell elongation as significantly different from the background. These were “Cell wall organisation”, “Phytohormone action” and “Cytoskeleton organisation”. The most specific significant subcategories within these were “pectin methylesterase” and “Fasciclin-type arabinogalactan protein”; “RALF/RALFL-precursor polypeptide”, “auxin-responsive genes *(SAUR)” and “transcriptional repressor *(Aux/IAA)”; and “alpha-beta-tubulin heterodimer”, respectively. For each of these, transcript levels were higher in the carpel 1 than carpel 2 samples, visible by a consistent shift in the ECDF plot to positive values (Figure 5A). This higher expression in carpel 1 was particularly evident when considering those members of each gene family with higher absolute expression levels (Figure 5B). As discussed below, the elevated expression of transcripts encoding pectin methylesterases, Fasciclin-type arabinogalactan proteins, SAUR and Aux/IAA proteins in carpel 1 is consistent with increased cell elongation driven by higher auxin levels in carpel 1 versus carpel 2. Our transcriptome analysis therefore suggests a molecular basis for the differential growth of the two adaxial carpels driving style deflection.

**Figure 5:**
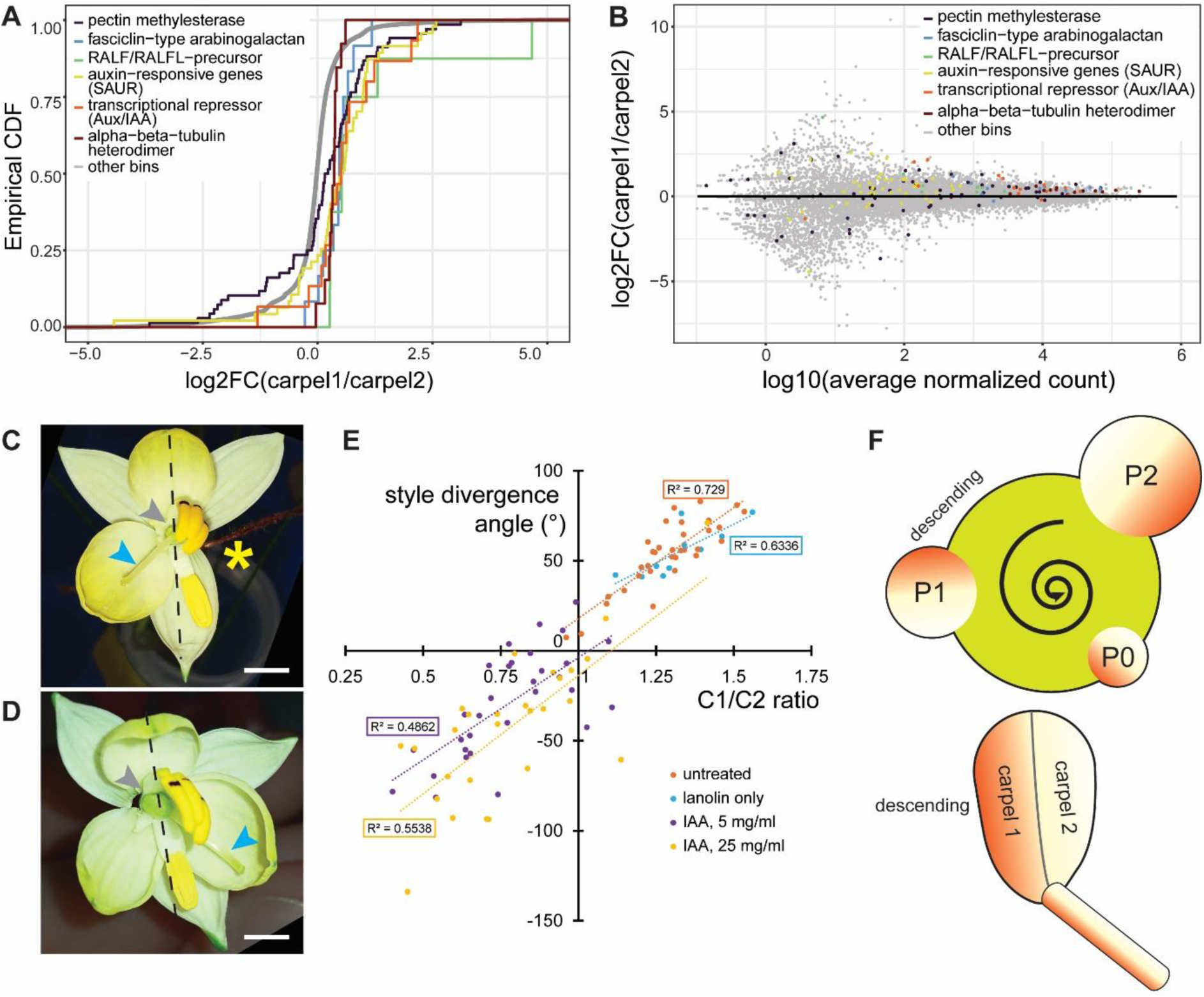
Differential auxin signaling between the two adaxial carpels underlies style deflection in *Cyanella alba* subsp*. Flavescens*. (**A**) Empirical cumulative distribution functions (ECDFs) of the log2 fold-changes (log2FC) in transcript abundance between carpels 1 and 2 are shown for selected Mercator bins with significantly different log2FC distributions compared to the transcriptome-wide background (‘other bins’). See Table S10 for results for all bins. (**B**) Log2 fold-changes (log2FC) in transcript abundance between carpels 1 and 2 are shown relative to the average base expression across all samples for all genes in the Mercator bins shown in (**A**). (**C**,**D**) *C. alba* flowers from left-biased plants after treatment with lanolin only (**C**) or lanolin with 5 mg/ml IAA (**D**) to carpel 2. Grey arrowheads indicate the side where two of the adaxial anthers were removed and lanolin was applied to the carpel 2. Blue arrowheads indicate styles. Dashed lines show the midline of the flower based on the adaxial petal and stamens. One of the three petals in (**C**) was removed (yellow asterisk). Scale bar is 5 mm. (**E**) Effect of auxin on style divergence angle and the ratio of carpel 1 to carpel 2 length (C1/C2 ratio). Positive divergence angles indicate style deflection away from carpel 1, negative angles towards carpel 1. Linear regression lines and R^2^ values are shown. Both IAA treated samples are significantly different from untreated and lanolin-only samples regarding both C1/C2 ratio and style divergence angle (p<0.001; t-test). (**F**) Schematic model for the development of a consistent floral handedness based on an initial asymmetry in auxin concentration in the initiating organs (gradients in circles, P2 to P0 from old to young; descending indicates the side of the primordium facing the next older primordium) at the shoot meristem (green). The direction of this initial asymmetry depends on the direction of the phyllotactic spiral (curved arrow, counter-clockwise in this illustration) and is later translated into style deflection (to the right in this illustration), by stronger elongation of carpel 1 than carpel 2.

### Auxin application to carpel 2 reverses the style orientation

To test the importance of differential auxin signaling between carpels 1 and 2 we asked whether external auxin application to carpel 2 could lead to straight styles or even a reversal of style orientation by causing carpel 2 to expand as much as, or more than, carpel 1. Lanolin with or without auxin was applied to the outside wall of carpel 2 in early-stage flowers (Figure 5C,D). After the flowers had opened, we dissected out the pistils and measured the lengths of carpels 1 and 2 along their midveins as well as the degree of style deflection relative to the long axis of the ovary. Across all treatment groups, there was a positive correlation between the ratio of carpel 1/carpel 2 lengths and the degree of style deflection; thus, the more uneven the expansion of the two carpels, the greater the deflection of the style is (Figure 5E; positive versus negative angles indicate styles deflecting away from carpel 1 versus towards carpel 1, respectively). However, while all untreated or lanolin-treated pistils had carpel 1/carpel 2 ratios above 1 and styles deflected away from carpel 1 (positive angles), application of auxin to carpel 2 shifted both measures towards lower values (*p*<0.001 for all comparisons of auxin treated to control groups based on *t*-test). This indicates that auxin application could equalize growth of the two carpels or even cause carpel 2 to be longer than carpel 1. As predicted, this led to the style being straight or even deflected away from carpel 2 and towards carpel 1. Together with the transcriptomic analysis, this indicates that in untreated plants higher auxin signaling in carpel 1 causes it to expand more than carpel 2, leading to style deflection away from carpel 1. Thus, unequal auxin-induced expansion of the carpels appears to be the primary cause of floral handedness of *C. alba* .

## DISCUSSION

Our results provide multiple lines of evidence indicating that *C. alba* possesses monomorphic rather than dimorphic enantiostyly. Molecular analyses revealed that the consistent handedness of the mirror-image flowers of individual plants within any single flowering season is not determined by their genotype. Rather, style deflection depends on the apparently random choice that each annually initiated shoot meristem makes concerning the handedness of the phyllotactic spiral. Once determined, the handedness of the phyllotactic spiral is stable throughout one growing season, which translates, albeit with occasional deviations, to a predictable and consistent direction of style and anther deflection per individual. We also show that this left-right asymmetric morphology has a simple cellular and developmental basis. Developmentally, style deflection is caused by differential cell elongation between the two adaxial carpels, coupled with a very stiff base to the ovary. The stronger elongation of one of the carpels results from increased auxin signaling and higher expression of cell-expansion promoting genes. Thus, our work identifies a non-genetic mechanism for ensuring consistent left-right handedness that links chirality across different levels of plant development to produce adaptive morphological variation.

The mechanism for determining stylar deflection based on the handedness of the phyllotactic spiral mirrors recent findings about leaf asymmetries in other plants. In tomato, *Arabidopsis thaliana* and several other species, the direction of more or less subtle LR asymmetries in leaves can be predicted from the handedness of the phyllotactic spiral ^11,16,43^. These asymmetries have been traced back to an unequal distribution of auxin and auxin signalling in initiating leaf primordia at the shoot apical meristem ^16^. In particular, the side of the primordium facing the next oldest (the so-called descending side) has higher auxin levels than the ascending side, resulting from the pattern of polar auxin transport. We therefore consider it highly likely that floral asymmetry in *C. alba* is ultimately caused by such an initial auxin asymmetry in initiating bract/floral primordia (Figure 5C) for two reasons. First, auxin maxima seem to determine the sites of organ initiation at the shoot apical meristem across all mono- and dicotyledonous plants studied, so also most likely in *C. alba*. Second, the asymmetry of auxin levels in initiating primordia in spiral phyllotaxis appears to result from the patterning mechanism of polar auxin transport itself that generates these auxin maxima, leading to what has been interpreted as an inherent mechanistic constraint on the starting conditions in an initiating primordium ^16^, which most likely applies to all species with spiral phyllotaxis.

This model raises three questions. How is the presumed initial auxin asymmetry translated into floral handedness? Is a similar mechanism operating in other plants with mirror-image flowers? What could be the functional significance of linking floral variation to the working of a core shoot-patterning mechanism? In a *C. alba* individual with counter-clockwise phyllotaxis and style deflection to the right, the left adaxial carpel (when viewed facing the flower) corresponds to the descending side of the initial primordium. Thus, according to our model the stronger auxin signalling on the descending side of the primordium ultimately triggers stronger cell expansion in the corresponding adaxial carpel (Figure 5C). When combined with a very stiff base of the ovary, such differential carpel elongation is predicted by our biophysical model to cause style deflection away from the descending side of the flower. Strikingly, our transcriptomic comparison between the more- and less-expanding adaxial carpels provides evidence for stronger auxin signalling in the carpel on the descending side of the initial primordium. This carpel has higher expression of Aux/IAA and SAUR encoding genes, both of which are induced by auxin signalling ^44,45^. The latter also have a well-established role in mediating acidification of the extracellular space, resulting in increased cell expansion as described by the acid-growth hypothesis ^44,46,47^.

In addition, the more strongly expanding adaxial carpel also shows higher expression of genes encoding extracellular pectin methylesterases and Fasciclin-like arabinogalactan proteins (FLAs). Overexpression of a pectin methylesterase in *A. thaliana* shoot meristems resulted in less rigid cell walls ^48^, presumably facilitating cell expansion, while overexpression of a FLA in cotton-fibre cells increased their elongation ^49^, suggesting that both activities also contribute to stronger expansion of the adaxial carpel on the descending side of the flower. Auxin and the auxin-response factor ARF3/ETTIN have been shown to induce pectin methylesterase activity or expression in tobacco and *A. thaliana*, respectively, suggesting a similar link in the descending carpel ^50–52^. At the same time, this carpel also shows higher expression of potential negative regulators of cell expansion, in particular Rapid Alkalinisation Factor (RALF) peptides and three pectin-methylesterase inhibitors, which limit cell expansion in *A. thaliana* roots and shoot meristems, respectively ^48,53^. RALF signalling has been shown to increase auxin biosynthesis in *A. thaliana* roots ^53^, providing another possible link between the differentially expressed gene categories in the *C. alba* carpels. The importance of differential auxin signalling between the two carpels is further supported by our external auxin application to carpel 2, which was sufficient to equalize growth between the two carpels or even lead to carpel 2 overgrowth, causing the style to become straight or deflected towards carpel 1.

Thus, even though the mechanistic details remain to be resolved, our findings suggest that increased auxin signalling in the more expanding adaxial carpel causes stronger cell elongation than in the less expanding one, leading to style deflection away from the more expanded side of the ovary. At the same time, it remains an open question how the information from the initial auxin asymmetry in the primordium could be maintained for the long time across many cell divisions between floral-meristem initiation and late stages of pistil morphogenesis.

An epigenetic process may be involved, but this remains speculative at present. However, this long duration between the presumed initial trigger and the later elaboration of the morphological asymmetry may explain the small fraction of ‘developmental deviations’ where individual flowers do not follow the pattern of style deflection predicted from the handedness of the phyllotactic spiral.

We propose that our model for *C. alba*, which resembles that proposed for leaves ^16^, may be more widely applicable to other species with floral asymmetry. An interaction between processes at the inflorescence meristem and flower development predicting the handedness of mirror-image flowers is also seen in other species with monomorphic enantiostyly, such as *Solanum rostratum* ^54^. The inflorescence of this species is a scorpioid monochasial cyme, i.e. each meristem terminates in a flower and inflorescence growth is continued by a single lateral meristem, whose position alternates relative to the respective main axis. Styles are almost always deflected towards the lateral branch, resulting in pendulum asymmetry ^55^, and suggesting that positional information from the primordium stage translates into floral handedness. A more directly comparable case is found in *Monochoria australasica* (also referred to as *Pontederia australasica*). This species of Australian perennial marsh plant shows inflorescence-level monomorphic enantiostyly, i.e. the flowers within one inflorescence have a consistent handedness, yet different inflorescences produced by the same individual can have a different floral handedness ^24^. Inspection of publicly available photographs of *M. australasica* inflorescences shows that the species has spiral phyllotaxis (Figure S10, Table S11). Across all available images where the handedness of both phyllotaxis and style deflection could be reliably assessed, we found that 7 of 9 inflorescences with a clockwise phyllotactic spiral had styles deflected to the right (one inflorescence mixed, one with L-flowers), and all 12 inflorescences with counter-clockwise phyllotaxis had styles deflected to the left. Thus, the handedness of the phyllotactic spiral again appears to determine floral handedness, although with an inverted relation compared to *C. alba*, indicating that the mechanism identified here also operates in other systems with monomorphic enantiostyly.

In plants with monomorphic enantiostyly where several flowers with different handedness are open on a given day, there is a risk of between-flower selfing (geitonogamy), as approximately half of the flowers should donate pollen efficiently to the other half ^21,27^. This could result in gamete wastage due to inbreeding depression and pollen discounting. Linking the handedness of individual flowers to phyllotaxis as a stable pattern-generating process at the shoot apical meristem can unify floral handedness across an inflorescence, thus limiting the mating costs of geitonogamy ^56^. At the same time, the random establishment of clockwise versus counter-clockwise phyllotaxis across individual shoot meristems results in an equal distribution of L- and R-biased inflorescences in a population, maximizing mating opportunities and effectively functioning like dimorphic enantiostyly. These beneficial effects would be maximal in a situation where the relation between phyllotactic and floral handedness was absolute, individuals only formed a single inflorescence, and there was a large floral display with many open flowers on a given day. The latter seems to be the case in *M. australasica*, yet its clonal growth with the possibility of forming more than one inflorescence per individual reintroduces some risk of within-plant selfing owing to between-inflorescence geitonogamy. In *C. alba*, each plant appears to form only a single inflorescence per growing season and daily floral display size is small, but often larger than one. This suggests that inflorescence-level uniformity of floral handedness can limit within-plant selfing and promote disassortative mating between individuals of opposite phyllotactic and floral handedness.

Thus, in conclusion we describe a conserved mechanism whereby plants exploit an inherent developmental constraint leading to chirality in a core developmental process to generate ecologically relevant morphological variation in the form of LR asymmetry.

## ACKNOWLEDGEMENTS

Plant material was collected under Cape Nature permit (CN35-87-25844). We are grateful to Marrietta and Barry Lubbe of Mertenhof farm and Barend Salomo and all members of the Wupperthal Original Rooibos Cooperative for permission and support in sampling *Cyanella alba*. We thank Sinéad Watchorn and Lia Hemerick for support in the field sampling. We thank Christian Kappel and Mathias Scharmann for suggestions on genome assembly and transcriptome analysis. Some computations were performed using facilities provided by the University of Cape Town’s ICTS High Performance Computing team. C.R. held a Harry Crossley Research Fellowship. M.K was funded by the Nederlandse Organisatie voor Wetenschappelijk Onderzoek (GSGT.2019.019). This work was supported by a grant from the Human Frontiers Science Program to E.D., N.I., S.C.H.B. and M.L. (grant number RGP0036/2021).

## DATA AVAILABILITY

Whole genome and transcriptome datasets generated in this work are available in NCBI under under BioProject PRJNA1123744.

## REFERENCES

1. Blum, M., Feistel, K., Thumberger, T. & Schweickert, A. The evolution and conservation of left-right patterning mechanisms. Development 141, 1603–1613 (2014).

2. Smyth, D. R. Helical growth in plant organs: mechanisms and significance. Development 143, 3272–3282 (2016).

3. Davison, A. et al. Formin Is associated with left-right asymmetry in the pond snail and the frog. Curr Biol 26, 654–660 (2016).

4. Kuroda, R., Endo, B., Abe, M. & Shimizu, M. Chiral blastomere arrangement dictates zygotic left-right asymmetry pathway in snails. Nature 462, 790–U112 (2009).

5. Nonaka, S., Shiratori, H., Saijoh, Y. & Hamada, H. Determination of left-right patterning of the mouse embryo by artificial nodal flow. Nature 418, 96–99 (2002).

6. Okada, Y., Takeda, S., Tanaka, Y., Belmonte, J. C. I. & Hirokawa, N. Mechanism of nodal flow: A conserved symmetry breaking event in left-right axis determination. Cell 121, 633–644 (2005).

7. Abe, M. & Kuroda, R. The development of CRISPR for a mollusc establishes the formin Lsdia1 as the long-sought gene for snail dextral/sinistral coiling. Development 146, (2019).

8. Grimes, D. T. & Burdine, R. D. Left-right patterning: Breaking symmetry to asymmetric morphogenesis. Trends Genet 33, 616–628 (2017).

9. Hirokawa, N., Tanaka, Y. & Okada, Y. Left–right determination: Involvement of molecular motor KIF3, cilia, and nodal flow. Cold Spring Harb. Perspect. Biol. 1, a000802 (2009).

10. Kuroda, R. A twisting story: how a single gene twists a snail? Mechanogenetics. Q. Rev. Biophys. 48, 445–452 (2015).

11. Korn, R. W. Anodic asymmetry of leaves and flowers and its relationship to phyllotaxis. Ann. Bot. 97, 1011–1015 (2006).

12. Allard, H. A. The ratios of clokwise and counterclockwise spirality observed in the phyllotaxy of some wild plants. Castanea 16, 1–6 (1951).

13. Sousa-Baena, M. S., Lohmann, L. G., Hernandes-Lopes, J. & Sinha, N. R. The molecular control of tendril development in angiosperms. New Phytol. 218, 944–958 (2018).

14. Buschmann, H. & Borchers, A. Handedness in plant cell expansion: a mutant perspective on helical growth. New Phytol (2019) doi:10.1111/nph.16034.

15. Reinhardt, D. et al. Regulation of phyllotaxis by polar auxin transport. Nature 426, 255– 60 (2003).

16. Chitwood, D. H. et al. Leaf asymmetry as a developmental constraint imposed by auxin-dependent phyllotactic patterning. Plant Cell 24, 2318–2327 (2012).

17. Gerbode, S. J., Puzey, J. R., McCormick, A. G. & Mahadevan, L. How the cucumber tendril coils and overwinds. Science 337, 1087–1091 (2012).

18. Thitamadee, S., Tuchihara, K. & Hashimoto, T. Microtubule basis for left-handed helical growth in Arabidopsis. Nature 417, 193–196 (2002).

19. Ishida, T., Kaneko, Y., Iwano, M. & Hashimoto, T. Helical microtubule arrays in a collection of twisting tubulin mutants of *Arabidopsis thaliana*. Proc. Natl. Acad. Sci. U. S. A. 104, 8544–8549 (2007).

20. Barrett, S. C. H. The evolution of plant sexual diversity. Nat Rev Genet 3, 274–84 (2002).

21. Jesson, L. K. & Barrett, S. C. H. Experimental tests of the function of mirror-image flowers. Biol. J. Linn. Soc. 85, 167–179 (2005).

22. Jesson, L. K. & Barrett, S. C. H. The genetics of mirror-image flowers. Proc. R. Soc. B-Biol. Sci. 269, 1835–1839 (2002).

23. Barrett, S. C. H. & Fairnie, A. L. M. The neglected floral polymorphism: mirror-image flowers emerge from the shadow of heterostyly. Evol. J. Linn. Soc. 3, kzae004 (2024).

24. Jesson, L. K. & Barrett, S. C. H. The comparative biology of mirror-image flowers. Int J Plant Sci 164, S237–S249 (2003).

25. Barrett, S. C. H. Darwin’s legacy: the forms, function and sexual diversity of flowers. Philos Trans R Soc Lond B Biol Sci 365, 351–68 (2010).

26. Barrett, S. C. H. Sexual interference of the floral kind. Heredity 88, 154–159 (2002).

27. Jesson, L. K. & Barrett, S. C. H. Enantiostyly: Solving the puzzle of mirror-image flowers. Nature 417, 707–707 (2002).

28. Minnaar, C. & Anderson, B. A combination of pollen mosaics on pollinators and floral handedness facilitates the increase of outcross pollen movement. Curr. Biol. CB 31, 3180–3184.e3 (2021).

29. Johnson, S. D., Midgley, J. J. & Illing, N. The enantiostylous floral polymorphism of *Barberetta aurea* (Haemodoraceae) facilitates wing pollination by syrphid flies. Ann. Bot. 132, 1107–1118 (2023).

30. Manning, J. C. & Goldblatt, P. A revision of Tecophilaeaceae subfam. Tecophilaeoideae in Africa. Bothalia Afr. Biodivers. Conserv. 42, (2012).

31. Paudel, B. R. et al. Loss of buzz pollination results in chronic pollen limitation in an enantiostylous plant. South Afr. J. Bot. 171, 592–601.

32. Dulberger, R. & Ornduff, R. Floral morphology and reproductive biology of four species of *Cyanella* (Tecophilaeaceae). New Phytol 86, 45- (1980).

33. Kappel, C., Huu, C. N. & Lenhard, M. A short story gets longer: recent insights into the molecular basis of heterostyly. J Exp Bot 68, 5719–5730 (2017).

34. Huu, C. N. et al. Presence versus absence of *CYP734A50* underlies the style-length dimorphism in primroses. Elife 5, e17956 (2016).

35. Shore, J. S. et al. The long and short of the S-locus in *Turnera* (Passifloraceae). New Phytol. 224, 1316–1329 (2019).

36. Mirabet, V., Besnard, F., Vernoux, T. & Boudaoud, A. Noise and robustness in phyllotaxis. PLOS Comput. Biol. 8, e1002389 (2012).

37. Marconi, M. & Wabnik, K. Shaping the organ: A biologist guide to quantitative models of plant morphogenesis. Front. Plant Sci. 12, (2021).

38. Waidmann, S. et al. Cytokinin functions as an asymmetric and anti-gravitropic signal in lateral roots. Nat. Commun. 10, 3540 (2019).

39. Nelder, J. A. & Mead, R. A simplex method for function minimization. Comput. J. 7, 308– 313 (1965).

40. Holland, M. A., Kosmata, T., Goriely, A. & Kuhl, E. On the mechanics of thin films and growing surfaces. Math. Mech. Solids MMS 18, 561–575 (2013).

41. Hamant, O. et al. Developmental patterning by mechanical signals in *Arabidopsis*. Science 322, 1650–5 (2008).

42. Bolger, M., Schwacke, R. & Usadel, B. MapMan visualization of RNA-seq data using Mercator4 functional annotations. in Solanum tuberosum: Methods and Protocols (eds. Dobnik, D., Gruden, K., Ramšak, Ž. & Coll, A.) 195–212 (Springer US, New York, NY, 2021). doi:10.1007/978-1-0716-1609-3_9.

43. Martinez, C. C., Chitwood, D. H., Smith, R. S. & Sinha, N. R. Left-right leaf asymmetry in decussate and distichous phyllotactic systems. Philos. Trans. R. Soc. Lond. B. Biol. Sci. 371, 20150412 (2016).

44. Du, M., Spalding, E. P. & Gray, W. M. Rapid auxin-mediated cell expansion. Annu. Rev. Plant Biol. 71, 379–402 (2020).

45. Krogan, N. T., Yin, X., Ckurshumova, W. & Berleth, T. Distinct subclades of Aux/IAA genes are direct targets of ARF5/MP transcriptional regulation. New Phytol. 204, 474– 483 (2014).

46. Spartz, A. K. et al. The SAUR19 subfamily of SMALL AUXIN UP RNA genes promote cell expansion. Plant J 70, 978–990 (2012).

47. Spartz, A. K. et al. SAUR inhibition of PP2C-D phosphatases activates plasma membrane H+-ATPases to promote cell expansion in *Arabidopsis*. Plant Cell 26, 2129– 2142 (2014).

48. Peaucelle, A. et al. Pectin-induced changes in cell wall mechanics underlie organ initiation in Arabidopsis. Curr Biol 21, 1720–6 (2011).

49. Huang, G.-Q. et al. A fasciclin-like arabinogalactan protein, GhFLA1, is involved in fiber initiation and elongation of cotton. Plant Physiol. 161, 1278–1290 (2013).

50. Bryan, W. H. & Newcomb, K. H. Stimulation of pectin methylesterase activity of cultured tobacco pith by indoleacetic acid. Physiol. Plant. 7, 290–297 (1954).

51. Andres-Robin, A. et al. Evidence for the regulation of gynoecium morphogenesis by ETTIN via cell wall dynamics. Plant Physiol. 178, 1222–1232 (2018).

52. Andres-Robin, A. et al. Immediate targets of ETTIN suggest a key role for pectin methylesterase inhibitors in the control of *Arabidopsis* gynecium development. Plant Signal. Behav. 15, 1771937.

53. Li, L. et al. RALF1 peptide triggers biphasic root growth inhibition upstream of auxin biosynthesis. Proc. Natl. Acad. Sci. U. S. A. 119, e2121058119 (2022).

54. Jesson, L. K., Kang, J., Wagner, S. L., Barrett, S. C. H. & Dengler, N. G. The development of enantiostyly. Am J Bot 90, 183–195 (2003).

55. Charlton, W. A. Pendulum symmetry. in Symmetry in Plants vol. Volume 4 61–87 (WORLD SCIENTIFIC, 1998).

56. Harder, L. D. & Barrett, S. C. H. Mating cost of large floral displays in hermaphrodite plants. Nature 373, 512–515 (1995).

