## Supporting Information Text, Figures, Tables, Methods for "Style deflection is determined by the handedness of phyllotaxis and auxin-induced differential cell elongation in a species with mirror-image flowers"

**Establishment of a reference genome assembly and sequencing of left- and right-biased individuals of *Cyanella alba* subsp. *flavescens***

To identify the presumed causal enantiostyly locus, we selected 25 *C. alba* individuals from sites in the Biedouw Valley, Western Cape, South Africa that had four or more open flowers of consistent handedness (15 left- and 10 right-handed plants, maximum flower number = 8; mean flower number = 5.2; Supp. Table S1) in September 2021. In addition, during the 2022 flowering season we selected 20 individuals (11 left- and 9 right-handed) which produced six or more flowers of consistent handedness (max = 12; mean = 7.5; Supp. Table S1) during a long-term monitoring study. Sixteen of the individuals from 2022 were derived from the same site in the Biedouw Valley as the plants sampled in 2021, while four plants were sampled from a site approximately 5 km North of the 2021 site. We subjected extracted DNA to individual Illumina whole-genome sequencing. Over 2 billion 150 bp paired end reads were produced across the 25 samples of the Illumina2021 dataset, with a median of 76.7 million reads produced per sample. Given an estimated haploid genome size of 0.98 Gb from flow cytometry, this represents roughly 23.5-fold coverage on average per sample. For the Illumina2022 set, sequencing of these 20 samples yielded an additional 1.6 billion 150 bp reads, with a median of 78.4 million reads produced per sample, representing roughly 24.3-fold coverage on average.

To facilitate the analysis of these sequences, we also established a reference genome assembly based on PacBio high-fidelity sequencing. For this we extracted DNA from an R-individual, C_209, which had formed seven consistently R-flowers in 2021. An R-morph was chosen, because an initial analysis of the Illumina2021 dataset using KmerGO2 (https://github.com/ChnMasterOG/KmerGO2) to identify morph-associated k-mers had indicated that in the 10 R-morph genomes the ratio of right- to left-associated k-mers was much higher than the converse ratio in the 15 L-morph genomes, suggesting that R-morphs are heterozygous for the causal locus and thus contain a high number of k-mers that are not found in L-morphs. Almost 21 Gb of reads were produced, which equates to a 21X genome coverage based on the flow cytometry genome size estimate, with a mean read length of 17,907 bp, and an N50 of 17,976 bp. Estimating the genome size based on k-mer frequencies using the counting tool KMC and the profiling tool GenomeScope2.0 resulted in an estimate of 700 Mb, thus smaller than the flow cytometry-based estimate of 0.98Gb. We used hifiasm to assemble two reference genomes, using either 1 Gb or 700 Mb as haploid genome sizes. BUSCO (Benchmarking Universal Single-Copy Orthologs) analysis indicated similarly high completeness for both assemblies. The “1 Gb” and “700 Mb” assemblies contain 2962 (91.6%)

and 2971 (91.9%) complete BUSCOs of the 3,236 BUSCO groups in the liliopsida_odb10 dataset. However, the former had a much higher percentage of duplicated BUSCOs (16.5%) than the latter (3.1%). Therefore, we continued our analysis with this latter assembly, which had a total length of 0.84 Gb (only slightly smaller than the flow cytometry-based estimate), and an N50 of 3.27 Mb.

**Testing for a bi-allelic locus determining floral handedness**

We tested two hypotheses regarding a possible simple, single-locus genetic basis of floral handedness in *C. alba*. The first was that one of the morphs is heterozygous and the other homozygous recessive. To test this, we performed a genome wide association study (GWAS) to identify single nucleotide polymorphisms (SNPs) associated with each phenotype. We aligned all 45 Illumina samples to the reference assembly and phenotype-associated SNPs were identified. The list of SNPs was filtered for sites where more than 80% of the individuals of one phenotype were heterozygous for the alternate alleles of the SNP, whereas in the other only <20% were heterozygous. We chose a relaxed threshold of 80% to account for the possibility that some individuals may have been mis-categorised, especially in the 2021 sample with its lower flower number per individual. In other words, we considered it possible that a genotypically L-plant might still have produced four R-flowers by a series of developmental errors or vice versa. The Illumina2021 and Illumina2022 datasets were first analysed separately, before combining all 45 samples (Table 1). In the Illumina2021 dataset, more heterozygous SNP sites were identified in R- than L-plants (Table 1), but the opposite pattern occurred in the Illumina2022 samples (Table 1). The combined dataset identified more SNPs that were heterozygous in L- than in R-plants. However, none of the identified SNPs were supported across more than one analysis. Therefore, we conclude that the identified SNPs are most likely unrelated to the putative *E* locus and that enantiostyly in *C. alba* does not appear to be governed by a bi-allelic Mendelian locus.

**Testing for a hemizygous locus determining floral handedness**

Our second hypothesis was that the putative *E* locus is hemizygous and present only in either L- or R-plants. As our reference genome was from a presumed R-plant, we first searched for an R-exclusive region by performing a coverage analysis. We calculated the number of Illumina reads mapping to each region of the haploid *C. alba* genome and compared left and right pools to identify regions where there was a significant difference in coverage. We used the two haploid assemblies generated by Hifiasm separately for this analysis. As above, we performed the analysis first with the Illumina2021 and Illumina2022 datasets separately, and then with the combined dataset of all 45 samples. Again, none of the windows identified as having significantly different coverage in L- compared to R-samples were shared between the

Illumina2021 and Illumina2022 datasets (Table S1), and no plausible candidate for a hemizygous region in the right-handed *C. alba* assembly was identified.

We next asked whether the putative *E* locus may be exclusively present in L-plants and thus undetectable with our R-plant reference assembly. Such a hemizygous region should be detectable based on L-plant exclusive *k*-mers in the L-pool. To search for L-morph exclusive k-mers, KmerGO2 version 2.01 was run on the Illumina2021 and Illumina2022 datasets sequentially, using the following settings: *k*-mer size: 40 bp, threads: 24 for kmc3 and filtering steps, eight threads for union step.

The Illumina2021 dataset showed the same result as obtained previously using 31-mers (Figure 1A). The number of right-associated *k*-mers (63,103) exceeds the number of left-associated *k*-mers (26,856). Two left-handed individuals (C_002 and C_010) show an unexpectedly high number of right-associated *k*-mers compared to other L-morphs; however, they still possess far fewer right-associated *k*-mers than the ten right-handed plants, and also possess more left-associated *k*-mers than all the R-morph plants, making it unlikely that they represent R-morph plants mis-categorised as L-morphs. In contrast with the Illumina2021 dataset, analysis of the Illumina2022 samples identified 443,456 left-, but only 73,343 right-associated *k*-mers. Surprisingly, all individuals, regardless of handedness, possessed more left- than right-associated *k*-mers. The much higher number of morph-associated kmers in the Illumina2022 than the Illumina2021 dataset likely reflects the smaller sample size of the former (20 as opposed to 25 individuals), and thus possibly higher rate of false-positive morph-associated k-mers. Among those morph-associated kmers, only 103 were shared by both datasets. Among them, 13 are left-associated in both datasets, 2 are right-associated in both datasets, 12 are left-associated in the Illumina2021 dataset but are right-associated in the Illumina 2022 dataset, and 76 are right-associated in the Illumina2021 dataset but are left-associated in the Illumina 2022 dataset. Since there is no agreement between the Illumina2021 and Illumina2022 datasets in this analysis, there is no strong support for a hemizygous region exclusive to left-handed plants.

**Testing for transcripts exclusively expressed in plants with one floral handedness**

As a complement to the coverage- or *k*-mer based search for a hemizygous region, we used RNA-seq of dissected styles (including the upper half of the ovary) to search for morph-specific transcripts. We harvested triplicate samples of styles from very early-, early- and mid-stage buds of plants with exclusively R- or L-flowers (see Supporting Information Text for definition of stages). No style deflection is visible in very early-stage buds, but this becomes detectable in early buds and obvious in mid-stage buds (see Figure 3A below). Thus, our samples should cover the critical time window when the orientation of style deflection is determined. We identified 137 differentially expressed genes between L- and R-samples (FDR: $\alpha < 0.05$), of which 52 were upregulated and 85 downregulated in R- versus L-buds (Table S2). None of the genes were exclusively expressed in L- or R-plants. Thus, the transcriptomic analysis does not provide evidence for genes only present or only expressed in one of the phenotypes. Taken together, our results show no convincing evidence for a simple genetic polymorphism determining floral handedness in *C. alba*.

### Biomechanical model

To describe the mechanical forces at play during ovary expansion, and to make predictions on what area in the ovary walls drive the expansion of the organ as a whole, we propose a two-fold apporach: first, based on the images of six ovaries of Cyanella alba flavescens (FigureS3A), we model them as polyhedron that best represent their morphological characteristics. This is achieved by considering as modelling polyhedron a skewed triangular prism topped by a truncated pyramid (Figures S4). We use a data-driven approach by approximating the two dimensional images of C. alba flavescens to the best fitting two dimensional projection of our polyhedron according to length, area, and perimeter of the image (Figure S5). Subsequently, under the hypothesis that differential wall expansion in C. alba flavescens results from the expansion of a carpel with the others being pulled by elastic coupling forces acting at a cellular level, we fit such a resulting polyhedron with another polyhedron with springs as edges and massless beads as nodes (Figure 3F). With numerical optimisation techniques like random search algorithms, we obtain the biomechanical parameters of the bead-spring system (i.e., the stiffness and elongation of the springs in the system) necessary for the latter to best reproduce the best fitting polyhedron identified by our data-driven approach (Figure S3B). The pipeline executing the analysis and parameter optimisation is written with Wolfram Mathematica, see our Mathematica notebook (Supplemental File lateR1.nb) for detail.

### Data-driven model: best fitting polyhedron to expanded ovary

#### A three-dimensional polyhedron to model *C. alba flavescens* ovaries

Based on images of *C. alba flavescens*, we modelled the ovaries at the early stage of development - i.e., before the direction of stylar deflection can be determined, as triangular prisms topped by truncated pyramids. These geometric solids effectively capture the main morphological features of the ovaries, including the base of the style, and the stylised curvature of the external walls (see Figure S4A,B). We made the reasonable assumption that the ovary has rotational symmetry of order 3 at this early developmental stage. Our analysis involved fitting the two-dimensional projection of an ovary from above (i.e., with vein 1 in the field-of-view) with the polyhedron that can best reproduce such a projection.

We labelled the nodes at the base of the style within the model polyhedron as $\alpha$, $\beta$, and $\gamma$. The nodes where the ovary roughly ceases to be vertically straight relative to the base of the ovary and begins to curve were denoted as $a$, $b$, and $c$ (see Figure S5A,B). We denoted the positions of the nodes of the polyhedron as $\mathbf{r}_j = (x_j, y_j, z_j)$, with $j = a, b, c, \alpha, \beta, \gamma$. In the microscopy images, nodes $c$ and $\gamma$ are not visible and, hence, their position could not be identified. Therefore, we assumed that $z_b = z_c$ and $z_\beta = z_\gamma$. This means that, in our models, carpel 2 and 3 were equivalent both before and after expansion. We set the base of the ovary at height $z = 0$. The outer edges of the solid trace the midveins of the three carpels, see Figure 4B for a section of the ovary overlapped with the top view of our model.

We denoted the length of the edges connecting the base of the ovary to $a$, $b$, and $c$ as $l_a = z_a$, $l_b = z_b$, $l_c = z_c$, respectively. Instead, we define the following lengths: $l_t = |\mathbf{r}_a - \mathbf{r}_b| = |\mathbf{r}_a - \mathbf{r}_c| = |\mathbf{r}_b - \mathbf{r}_c|$, $l_s = |\mathbf{r}_\alpha - \mathbf{r}_\beta| = |\mathbf{r}_\alpha - \mathbf{r}_\gamma| = |\mathbf{r}_\beta - \mathbf{r}_\gamma|$, and $l_u = |\mathbf{r}_a - \mathbf{r}_\alpha| = |\mathbf{r}_b - \mathbf{r}_\beta| = |\mathbf{r}_c - \mathbf{r}_\gamma|$. Note that

$$l_t = \frac{2}{\sqrt{3}}|x_a - x_b|. \tag{S1}$$

$$l_s = \frac{2}{\sqrt{3}}|x_\alpha - x_\beta|, \tag{S2}$$

$$l_u = \sqrt{(z_a - z_\alpha)^2 + \left(\frac{l_t - l_s}{\sqrt{3}}\right)^2}. \tag{S3}$$

Finally, though not part of the polyhedron, we denoted the shortest length of the segments connecting nodes $\alpha$, $\beta$, and $\gamma$ with the base of the ovary as $l_\alpha = z_\alpha$, $l_\beta = z_\beta$, and $l_\gamma = z_\gamma$, see Figure S4A for a schematic representation.

**Morphological features of expanded ovaries**

We used the same solid to model ovaries in the late stage of development, except in the latter case the solid was skewed. We experimentally observe that stylar bending is driven by differential carpel elongation (see Result section in the main text), with the arbitrarily denominated carpel 1 being the most elongating one. The inner and outer walls of carpel 1 are, without loss of generality from now on considered as those connecting the base of the ovary with nodes $a$ and $\alpha$. We made the assumption that expansion of carpels 2 and 3 is comparable. Therefore, we maintained the assumption that $z_b = z_c$ and $z_\beta = z_\gamma$ even after ovary expansion.

Morphological features of six images of the lateral view of ovaries of *C. alba flavescens* were measured with Wolfram Mathematica, see our Mathematica notebook (Supplemental File lateR1.nb) for details. The six images (Figure S3) consist of two ovaries with left-handed symmetry (labelled by L1 and L2) and three with right-handed symmetry (R1, R2, and R3) at late developmental stage, and one right-handed ovary at mid developmental stage (R4). Specifically, we measured the distance between the base of the ovary and the base of the style, i.e., $\overline{z}_\alpha$ and $\overline{z}_\beta$, the perimeter of the ovary $\overline{P}$, its area $\overline{A}$, the length of the ovary base $|\overline{x}_a - \overline{x}_b|$, and the length of the style base $|\overline{\mathbf{r}}_\alpha - \overline{\mathbf{r}}_\beta|$, see Figure S5C and D. Measured quantities are listed in Table S7.

Table S7: Morphological measurements. All numerical values except for $\overline{A}$ are expressed in millimeters. $\overline{A}$ is expressed in millimeters squared.

| Ovary ID | $\overline{z}_\alpha$ | $\overline{z}_\beta$ | $\overline{P}$ | $\overline{A}$ | $\lvert\overline{x}_a - \overline{x}_b\rvert$ | $\lvert\overline{\mathbf{r}}_\alpha - \overline{\mathbf{r}}_\beta\rvert$ |
|---|---|---|---|---|---|---|
| L1 | 5.3 | 4.7 | 14.0 | 11.6 | 2.5 | 0.8 |
| L2 | 4.1 | 3.9 | 11.9 | 8.8 | 2.4 | 0.7 |
| R1 | 4.1 | 3.7 | 12.1 | 9.7 | 2.8 | 0.6 |
| R2 | 4.2 | 3.6 | 12.4 | 9.8 | 2.4 | 0.7 |
| R3 | 3.3 | 2.8 | 9.5 | 6.0 | 2.0 | 0.6 |
| R4 | 2.2 | 2.0 | 6.5 | 2.7 | 1.4 | 0.5 |

**Two-dimensional projection and fitting**

We projected the expanded polyhedron defined before on the vertical plane passing through the nodes $a$ and $\alpha$ and perpendicular to the edges that connect $b$ and $c$, see Figure S4C,D. With simple trigonometry, it is possible to show that the equation for such a projection is

$$\begin{aligned} T(x) = &\left[z_a + \frac{(z_\alpha - z_a)}{\frac{l_t - l_s}{\sqrt{3}}}x\right]\Theta\left(\frac{l_t - l_s}{\sqrt{3}} - x\right) \\ &+ \left[z_\alpha + (z_\beta - z_\alpha)\frac{x - \frac{l_t-l_s}{\sqrt{3}}}{\frac{\sqrt{3}}{2}l_s}\right]\Theta\left(-\frac{l_t - l_s}{\sqrt{3}} + x\right)\Theta\left(\frac{l_t - l_s}{\sqrt{3}} + \frac{\sqrt{3}}{2}l_s - x\right) \\ &+ \left[z_\beta + (z_b - z_\beta)\frac{x - \frac{l_t-l_s}{\sqrt{3}} - \frac{\sqrt{3}}{2}l_s}{l_t - \frac{l_t-l_s}{\sqrt{3}} - \frac{\sqrt{3}}{2}l_s}\right]\Theta\left(-\frac{l_t - l_s}{\sqrt{3}} - \frac{\sqrt{3}}{2}l_s + x\right)\Theta\left(\frac{\sqrt{3}}{2}l_s - x\right), \end{aligned} \tag{S4}$$

where $\Theta(\cdot)$ is the Heaviside theta function. The area $A$ of the projected polyhedron can be calculated using the previous equation as functions of $z_a$, $z_b$, $z_\alpha$, and $z_\beta$:

$$A(z_a, z_b, z_\alpha, z_\beta) = \int_0^\infty \mathrm{d}x\, T(x), \tag{S5}$$

The perimeter of the projected polyhedron, instead, is

$$P(z_a, z_b, z_\alpha, z_\beta) = z_a + |\mathbf{r}_a - \mathbf{r}_\alpha| + |\mathbf{r}_\alpha - \mathbf{r}_\beta| + |\mathbf{r}_\beta - \mathbf{r}_b| + z_b + |x_a - z_b|. \tag{S6}$$

This projection of the polyhedron wass then used to fit the two dimensional images of *C. alba flavescens* through optimisation of the model quantities $z_a, z_b, z_\alpha, z_\beta$ to best reproduce the morphological parameters of the six analysed images of *C. alba flavescens* according to the target function

$$f_D(z_a, z_b, z_\alpha, z_\beta) = (z_\alpha - \overline{z}_\alpha)^2 + (z_\beta - \overline{z}_\beta)^2 + \left(P(z_a, z_b, z_\alpha, z_\beta) - \overline{P}\right)^2 + \left(A(z_a, z_b, z_\alpha, z_\beta) - \overline{A}\right)^2 . \tag{S7}$$

We minimised this function by Nelder Mead method (Nelder & Mead, 1965) of the function NMinimize of Wolfram Mathematica to obtain the optimal values for $z_a, z_b, z_\alpha, z_\beta$, and we denoted them as $z_j^{\text{min}}$. Such optimal values are listed in Table S8

Table S8: Numerical values for best fitting polyhedra. Numerical values are expressed in millimeters.

| Ovary ID | $z_a^{\text{min}}$ | $z_\alpha^{\text{min}}$ | $z_\beta^{\text{min}}$ | $z_b^{\text{min}}$ |
|---|---|---|---|---|
| L1 | 4.1 | 5.3 | 3.2 | 4.7 |
| L2 | 3.3 | 4.1 | 1.0 | 3.4 |
| R1 | 2.9 | 4.1 | 2.6 | 3.7 |
| R2 | 3.7 | 4.2 | 2.6 | 3.6 |
| R3 | 2.3 | 3.3 | 2.0 | 2.8 |
| R4 | 1.5 | 2.2 | 1.5 | 2.0 |

### Biomechanical model: a bead-spring system to reproduce ovary expansion

In this section, we modelled the family of polyhedra presented in the previous section as systems of beads and springs, with springs representing the edges and beads representing the nodes of a polyhedron (Figure 3F). With our bead-spring system, we are interested in reproducing the polyhedra identified in the previous section as the best fitting to the expanded ovaries of *C. alba flavescens*. As such, we are interested in a static configuration of such a bead-spring system, rather than in an oscillatory behaviour.

**Initial conditions before ovary expansion**

In principle, to model the expansion of an ovary, it is necessary to observe both the initial morphology at an early stage of development, i.e., before stylar handedness could be determined, and the final morphology, after stylar deflection and differential ovary expansion have occurred. More specifically, it is necessary to know the equilibrium length of the springs of the system at the beginning of the expansion process. Under the assumption that ovary expansion only occurs along the $z$-axis, morphological features such as length of the style base $|\mathbf{r}_\alpha - \mathbf{r}_\beta|$, length of the base of an ovary $|x_a - x_b|$, and their derived quantities $l_t$ and $l_s$ can be considered constant throughout the expansion process, and we used the values reported in Table S7. Similarly, as we assumed that expansion only occurs in the first carpel, $l_b$, $l_\beta$, and $l_u$ can be considered constant. Finally, due to the assumption of rotational symmetry of order 3, it follows that, before ovary expansion begins, $l_a = l_b$ and $l_\alpha = l_\beta$.

Unfortunately, we could only analyze the morphological features of one ovary at early stage development (Figure S5A,B), which we used to infer $l_b$ and $l_\beta$ (and, therefore, their depending quantity $l_u$) of the expanded ovaries. Based on the analysis of Figure S5A and B, we measured five morphological characteristics: the distances between base of the ovary and nodes $a$, $\alpha$, $\beta$, and $b$ ($z_a^{\text{early}}$, $z_b^{\text{early}}$, $z_\alpha^{\text{early}}$, and $z_\beta^{\text{early}}$, respectively), and the length of the ovary base ($|x_a^{\text{early}} - x_b^{\text{early}}|$). The morphological features are listed in Table S9. Then, we defined the equilibrium length of the

Table S9: Morphological measurements of the ovary in Figure S5A,B at early stage development. Numerical values are expressed in millimeters.

| $z_a^{\text{early}}$ | $z_b^{\text{early}}$ | $z_\alpha^{\text{early}}$ | $z_\beta^{\text{early}}$ | $\lvert x_a^{\text{early}} - x_b^{\text{early}}\rvert$ |
|---|---|---|---|---|
| 0.7 | 0.5 | 1.6 | 1.6 | 1.3 |

springs $a$ and $b$ of the ovary in Figure S5A,B as the average between the two length measurements, i.e., $l_b = (\overline{z}_a + \overline{z}_b)/2 = 0.6$mm. Similarly, we defined $l_\beta = (\overline{z}_\alpha + \overline{z}_\beta)/2 = 1.6$mm.

We used the length of the ovary base of the images of expanded ovaries (Figure S3) in relation to the same quantity in the ovary at an early stage (Figure S5A,B) to infer $l_b$ and $l_\beta$ for the expanded ovaries. To do so, we defined the scaling factor

$$\text{Scaling factor} = \frac{|x_a - x_b|}{|x_a^{\text{early}} - x_b^{\text{early}}|}, \tag{S8}$$

to re-scale $l_b$ and $l_\beta$ according to the following equations:

$$l_b = \frac{z_a^{\text{early}} + z_b^{\text{early}}}{2} \frac{|x_a - x_b|}{|x_a^{\text{early}} - x_b^{\text{early}}|}, \tag{S9a}$$

$$l_\beta = \frac{z_\alpha^{\text{early}} + z_\beta^{\text{early}}}{2} \frac{|x_a - x_b|}{|x_a^{\text{early}} - x_b^{\text{early}}|}. \tag{S9b}$$

The scaling factors are listed in Table S10.

Table S10: Scaling factors

| Ovary ID | L1 | L2 | R1 | R2 | R3 | R4 |
|---|---|---|---|---|---|---|
| Scaling factor | 2.0 | 2.1 | 2.2 | 2.1 | 1.7 | 1.1 |

**Bead-spring system**

Before expansion, the system consisted of a regular prism with triangular basis topped by a truncated pyramid. The vertices of this polyhedron are represented by beads of mass $m$, whereas the edges are replaced by spring with equilibrium length $l_j$, with $j = a, b, c, \alpha, \beta, \gamma$, and spring constant as follows. We made the assumption that corresponding walls of different carpels possessed the same stiffness. Therefore, external springs, corresponding to springs denoted by $a, b, c$, have spring constant $k_e$, which is also the elastic constant of spring connecting $a$ to $\alpha$, $b$ to $\beta$, and $c$ to $\gamma$. Inner springs, i.e., springs denoted by $\alpha, \beta, \gamma$ have elastic constant $k_i$. Springs connecting $a, b, c$ have elastic constant $k_u$, whereas springs connecting $\alpha, \beta, \gamma$ - i.e., the base of the style, have elastic constant $k_s$.

**Dynamic equations**

Typically, the static equilibrium configuration $\mathbf{z}^{\text{eq}} = (z_a^{\text{eq}}, z_b^{\text{eq}}, z_c^{\text{eq}}, z_\alpha^{\text{eq}}, z_\beta^{\text{eq}}, z_\gamma^{\text{eq}})$ of positions of the beads consists in minimizing the potential energy of the system. In our case, directly solving for the equilibrium configuration was computationally demanding. Therefore, we tackled this challenge by solving the dynamic equations controlling the system. Through iteration of the dynamic solution, we identified an approximate static solution, thereby increasing the computational efficiency.

The dynamics of the bead-spring system is described by a system of second order linear ODEs. Given that our goal was to find the equilibrium configuration $\mathbf{z}^{\mathrm{eq}}$, we assumed, without loss of generality, that all beads in the system have identical mass. Consequently, we treat the elastic constants, denoted by $k$, as per unit mass. This assumption is justified as our model neglects gravitational and non-conservative forces, indicating that the equilibrium configuration - defined as the configuration that minimizes potential energy, is independent of the mass of the beads. Then, the dynamics equations are

$$\begin{cases} \ddot{\mathbf{r}}_a = -k_e(\mathbf{r}_a - l_a\mathbf{v}) - k_t(\mathbf{r}_a - \mathbf{r}_b - \mathbf{l}_{ab}) - k_t(\mathbf{r}_a - \mathbf{r}_c - \mathbf{l}_{ac}) - k_u(\mathbf{r}_a - \mathbf{r}_\alpha - \mathbf{l}_{a\alpha}), \\ \ddot{\mathbf{r}}_b = -k_e(\mathbf{r}_b - l_b\mathbf{v}) - k_t(\mathbf{r}_b - \mathbf{r}_a - \mathbf{l}_{ba}) - k_t(\mathbf{r}_b - \mathbf{r}_c - \mathbf{l}_{bc}) - k_u(\mathbf{r}_b - \mathbf{r}_\beta - \mathbf{l}_{b\beta}), \\ \ddot{\mathbf{r}}_c = -k_e(\mathbf{r}_c - l_c\mathbf{v}) - k_t(\mathbf{r}_c - \mathbf{r}_a - \mathbf{l}_{ca}) - k_t(\mathbf{r}_c - \mathbf{r}_b - \mathbf{l}_{cb}) - k_u(\mathbf{r}_c - \mathbf{r}_\gamma - \mathbf{l}_{c\gamma}), \\ \ddot{\mathbf{r}}_\alpha = -k_i(\mathbf{r}_\alpha - l_\alpha\mathbf{v}) - k_s(\mathbf{r}_\alpha - \mathbf{r}_\beta - \mathbf{l}_{\alpha\beta}) - k_s(\mathbf{r}_\alpha - \mathbf{r}_\gamma - \mathbf{l}_{\alpha\gamma}) - k_u(\mathbf{r}_\alpha - \mathbf{r}_a - \mathbf{l}_{\alpha a}), \\ \ddot{\mathbf{r}}_\beta = -k_i(\mathbf{r}_\beta - l_\beta\mathbf{v}) - k_s(\mathbf{r}_\beta - \mathbf{r}_\alpha - \mathbf{l}_{\beta\alpha}) - k_s(\mathbf{r}_\beta - \mathbf{r}_\gamma - \mathbf{l}_{\beta\gamma}) - k_u(\mathbf{r}_\beta - \mathbf{r}_b - \mathbf{l}_{\beta b}), \\ \ddot{\mathbf{r}}_\gamma = -k_i(\mathbf{r}_\gamma - l_\gamma\mathbf{v}) - k_s(\mathbf{r}_\gamma - \mathbf{r}_\alpha - \mathbf{l}_{\gamma\alpha}) - k_s(\mathbf{r}_\gamma - \mathbf{r}_\beta - \mathbf{l}_{\gamma\beta}) - k_u(\mathbf{r}_\gamma - \mathbf{r}_c - \mathbf{l}_{\gamma c}), \end{cases} \tag{S10}$$

where $\mathbf{v} = (0, 0, 1)$ is a unit vector in the $z$-direction, and $\mathbf{l}_{ij}$ is a vector with origin in $\mathbf{r}_i$ and directed towards $\mathbf{r}_j$ with length $l_{ij}$. Given that the nodes were constrained to oscillate in the $z$-direction, our system of $18$ differential equations reduced to $6$ differential equation for the $z$-component only of the position vector **r** of the nodes. With simple trigonometry, it is possible to show that

$$(\ddot{\mathbf{r}}_j)_z = \ddot{z}_j, \tag{S11a}$$

$$(\mathbf{r}_j - l_j\mathbf{v})_z = z_j - l_j, \tag{S11b}$$

$$(\mathbf{r}_j - \mathbf{r}_m - \mathbf{l}_{jm})_z = \begin{cases} (z_j - z_m)\left(1 - \frac{l_t}{\sqrt{l_t^2 + (z_j - z_m)^2}}\right) & \text{if} \quad j, m = a, b, c, \\ (z_j - z_m)\left(1 - \frac{l_s}{\sqrt{l_s^2 + (z_j - z_m)^2}}\right) & \text{if} \quad j, m = \alpha, \beta, \gamma, \\ (z_j - z_m)\left(1 - \frac{l_u}{\sqrt{\frac{(l_t - l_s)^2}{3} + (z_j - z_m)^2}}\right) & \text{if} \quad (j, m) = (a, \alpha), (b, \beta), (c, \gamma). \end{cases} \tag{S11c}$$

As we were interested in the static solution, we used the following initial conditions for every bead $j$:

$$\begin{cases} \ddot{z}_j = 0, \\ \dot{z}_j = 0. \end{cases} \tag{S12}$$

**Static solution**

We were interested in finding the equilibrium configuration of the system, i.e., the initial conditions $z_j(t = 0) = z_j^0 = z_j^{\text{eq}}$ such that $\dot{z}_j(t)|_{z_j^{\text{eq}}} = 0$ and $\ddot{z}_j(t)|_{z_j^{\text{eq}}} = 0$ at every time $t$. We proceeded by iterations, i.e., we subsequently solved the system of Equations (S10) with initial conditions $z_j^0[n] = z_j^{\text{eq}}[n-1]$ and $\dot{z}_j^0[n] = 0$, where $n$ indicates the $n$th iteration of the solving process, and where $z_j^0 = l_j$. We defined the equilibrium position of the node $j$ as

$$z_j^{\text{eq}} = \lim_{n\to\infty} z_j^{\text{eq}}[n]. \tag{S13}$$

The solutions of Equations (S10) are oscillatory (Figure S5) and, as such, $z_j[n] = z_j[n](t)$. Therefore, we defined the equilibrium conditions as the time average of such quantities, i.e.,

$$z_j^{\text{eq}}[n] = \lim_{T\to\infty} \frac{1}{T} \int_0^T \mathrm{d}t\, z_j[n](t). \tag{S14}$$

In our analyses, we used a more practical approach by integrating between $t = 0$ and $t = 200$ sec, and we considered $z_j$ to be at their equilibrium after the iteration $n = N$ if, for every $j =$

$a, b, c, \alpha, \beta, \gamma,$

$$\frac{\left|z_j^{\text{eq}}[N] - z_j^{\text{eq}}[N-1]\right|}{z_j^{\text{eq}}[N-1]} < 0.0001. \tag{S15}$$

Then, in our analysis, we approximated $z_j^{\text{eq}} \simeq z_j^{\text{eq}}[N]$.

### Optimisation: linking mechanical and data-driven models

We used the procedure as described in the previous section to find the polyhedron resulting from bead-spring model under static solution that best reproduced the polyhedron as identified by our data-driven modelling in the first section. To do so, we minimized the function $f_M$ defined by

$$f_M\left(l_a^{\text{opt}}, l_\alpha^{\text{opt}}, l_\beta^{\text{opt}}, l_b^{\text{opt}}, k_e^{\text{opt}}, k_i^{\text{opt}}, k_s^{\text{opt}}, k_t^{\text{opt}}\right) = \left(z_a^{\text{eq}} - z_a^{\text{min}}\right)^2 + \left(z_b^{\text{eq}} - z_b^{\text{min}}\right)^2 + \left(z_\alpha^{\text{eq}} - z_\alpha^{\text{min}}\right)^2 + \left(z_\beta^{\text{eq}} - z_\beta^{\text{min}}\right)^2. \tag{S16}$$

Parameters $z_j^{\text{eq}}$ are actually functions of the dynamic parameters of the models, i.e., $l_a$, $l_b$, $l_\alpha$, $l_\beta$, and $k_e$, $k_i$, $k_s$, $k_t$, $k_u$. In sticking with the assumption that only $l_a$ and $l_\alpha$ can elongate unconstrained with respect of the initial length of early stage ovaries, we allowed for a maximum elongation of $l_b$ and $l_\beta$ of $10$% respect to their initial length.

In our optimization, we used the measurements reported in Table S8 for $z_j^{\text{min}}$ and their derived quantities $l_s$, $l_t$, and $l_u$. Instead, $l_b$ and $l_\beta$ have been inferred according to the procedure described in the section "Initial conditions before ovary expansion" and the re-scaling factors listed in Table S10.

We performed parameter optimisation with the function NMinimize in Mathematica, by using the Nelder-Mead controlled random search algorithm. The results of our parameter optimisation are listed in Table S11. We used such parameters to reconstruct, through our mechanistic model and Equation (S4), the equilibrium configuration of the bead-spring system that best reproduced the best fitting polyhedron after geometrical optimisation. Figure S3B and Table S12 show, respectively, the resulting fitting polyhedron, and a summary of the change in morphological characteristics of the six ovaries of *Cyanella alba flavescens* that we analysed.

Table S11: Numerical values for optimised parameters of the biomechanical model. Lengths are measured in millimeters, spring constants in Newton per millimeter per kilogram.

| Ovary ID | $l_a^{\text{opt}}$ | $l_\alpha^{\text{opt}}$ | $l_\beta^{\text{opt}}$ | $l_b^{\text{opt}}$ | $k_e^{\text{opt}}$ | $k_i^{\text{opt}}$ | $k_s^{\text{opt}}$ | $k_t^{\text{opt}}$ |
|---|---|---|---|---|---|---|---|---|
| L1 | 5.6 | 6.3 | 3.5 | 1.2 | 7.3 | 1.3 | 16.2 | 9.8 |
| L2 | 3.8 | 4.1 | 3.2 | 1.2 | 9.5 | 9.8 | 16.9 | 1.1 |
| R1 | 3.1 | 4.1 | 3.3 | 1.5 | 9.3 | 1.0 | 15.6 | 2.1 |
| R2 | 4.9 | 4.8 | 3.1 | 1.4 | 3.3 | 8.5 | 18.8 | 8.9 |
| R3 | 2.9 | 3.5 | 2.6 | 1.2 | 9.8 | 10.0 | 19.2 | 9.8 |
| R4 | 1.7 | 2.2 | 0.8 | 1.8 | 8.2 | 1.9 | 16.7 | 3.3 |

Table S12: Elongation of polyhedron edges compared to equilibrium length of the respective representing spring in our biomechanical models. Positive values correspond to edges which are shorter than the respective equilibrium length and are, therefore, compressed. Similarly, negative values correspond to edges that are longer than the respective equilibrium length and are, therefore, stretched. Numerical values are expressed in millimeters.

| Ovary ID | $l_a^{\text{opt}} - z_a$ | $l_\alpha^{\text{opt}} - z_\alpha$ | $l_\beta^{\text{opt}} - z_\beta$ | $l_b^{\text{opt}} - z_b$ |
|---|---|---|---|---|
| L1 | 1.2 | 1.1 | -0.4 | -0.8 |
| L2 | 0.5 | 0.1 | -0.2 | -0.1 |
| R1 | 0.3 | 0.1 | -0.3 | -0.2 |
| R2 | 1.4 | 0.5 | -0.4 | -0.4 |
| R3 | 0.4 | 0.2 | -0.2 | -0.1 |
| R4 | 0.2 | 0.0 | -0.1 | -0.1 |

**Supporting Information Figures**

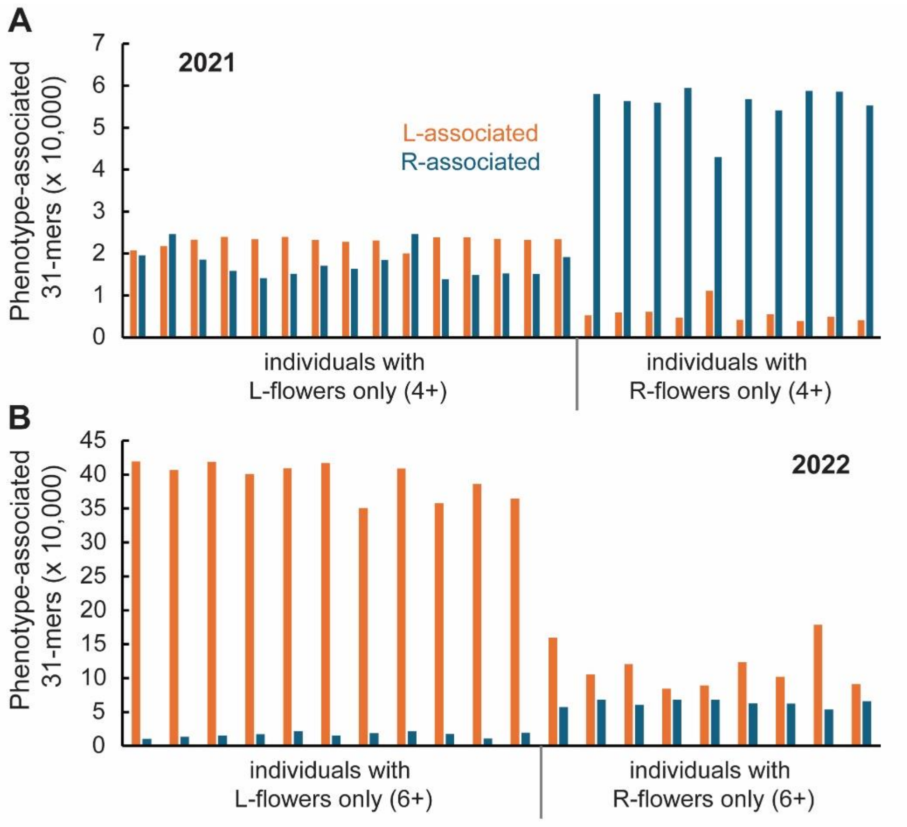

**Figure S1: Lack of evidence for a genomic region associated with style orientation**

(**A**,**B**) Phenotype-associated *k*-mer composition of *C. alba* individuals in the Illumina2021 (**A**) and Illumina2022 (**B**) datasets. *K*-mers associated with either the L- or R-pools were identified using KmerGO2. The total number of L- and R-associated 31-mers in each individual was calculated and plotted.

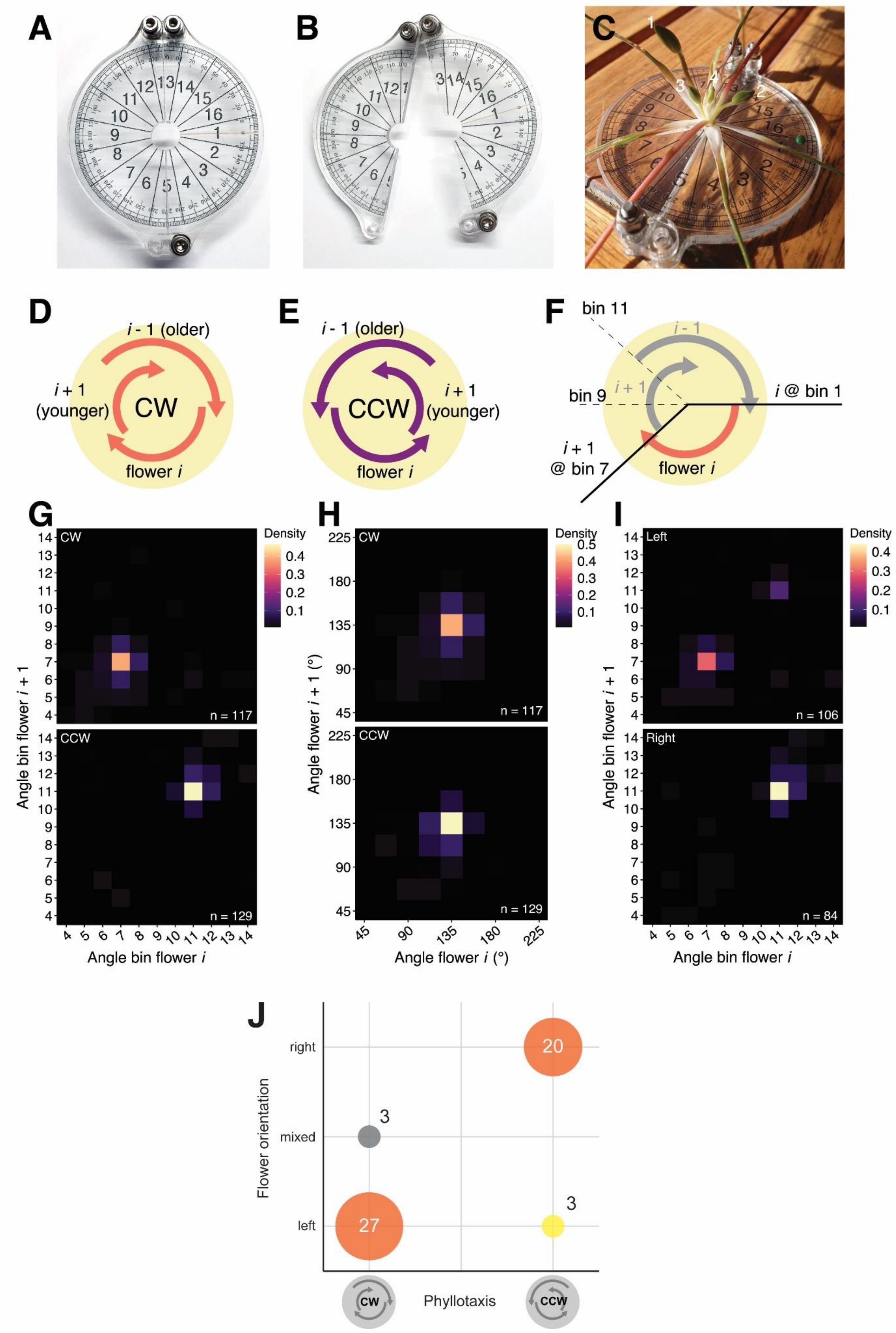

**Figure S2: *Cyanella alba* exhibits spiral phyllotaxis.**

(**A**, **B**) Overview of the hinged protractor for phyllotactic measurements in closed (**A**) and open (**B**) conformation. Multiple diameters of the central hole were produced and used, as a snug fit provides the most accurate measurements.

(**C**) Protractor around a *C. alba* plant exhibiting CW spiral phyllotaxis. Flower buds are labelled 1-4 (oldest to youngest). Two stalks of open flowers are also visible.

(**D**, **E**) Schematic diagram of three flowers on a plant with CW (**D**) or CCW (**E**) phyllotaxis.

(**F**) Visualization of phyllotactic measurements of a flower i in relation to the next (younger) flower i + 1 on a plant with CW phyllotaxis. In this scenario, the assigned angle bin would be '7'.

(**G**) 2D heatmap of measured angle bins of flowers i against i + 1 for all analyzed *C. alba* plants, separated based on the chirality of their phyllotaxis. Colour-scaled density represents the proportion of angles per panel falling in a specific 2D bin.

(**H**) Translation of the bins in (**G**) to degrees.

(**I**) 2D heatmap of angle bins separated on plants based on their predominant stylar orientation (left or right).

(**J**) Correlation of phyllotactic direction (CW: clockwise, CCW: counter-clockwise) and floral handedness of plants analyzed in 2024. Numbers in the diagram indicate the number of individual plants. The determination of floral handedness is based on between 2 and 5 open flowers and sufficiently old buds per plant.

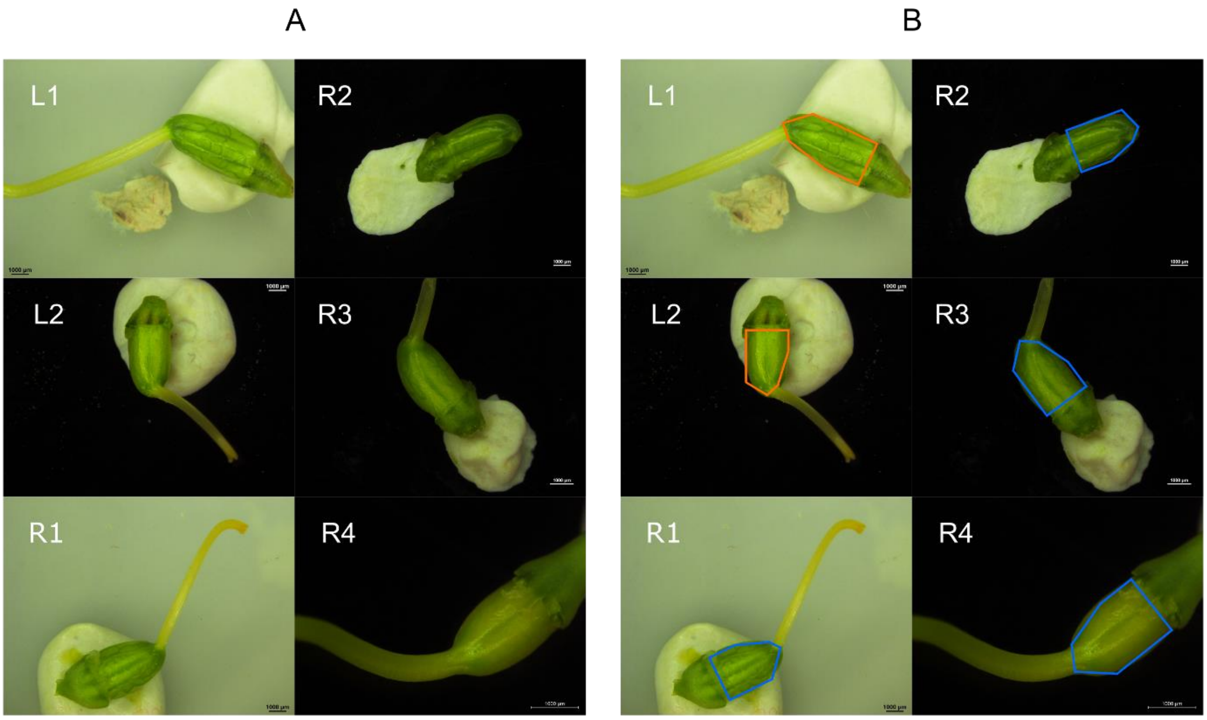

**Figure S3: Starting images for gynoecium modelling.**

(**A**) Lateral images of six ovaries of *C. alba flavescens*. (**B**) The same images are overlaid with the respective fitting polyhedron after parameter optimisation.

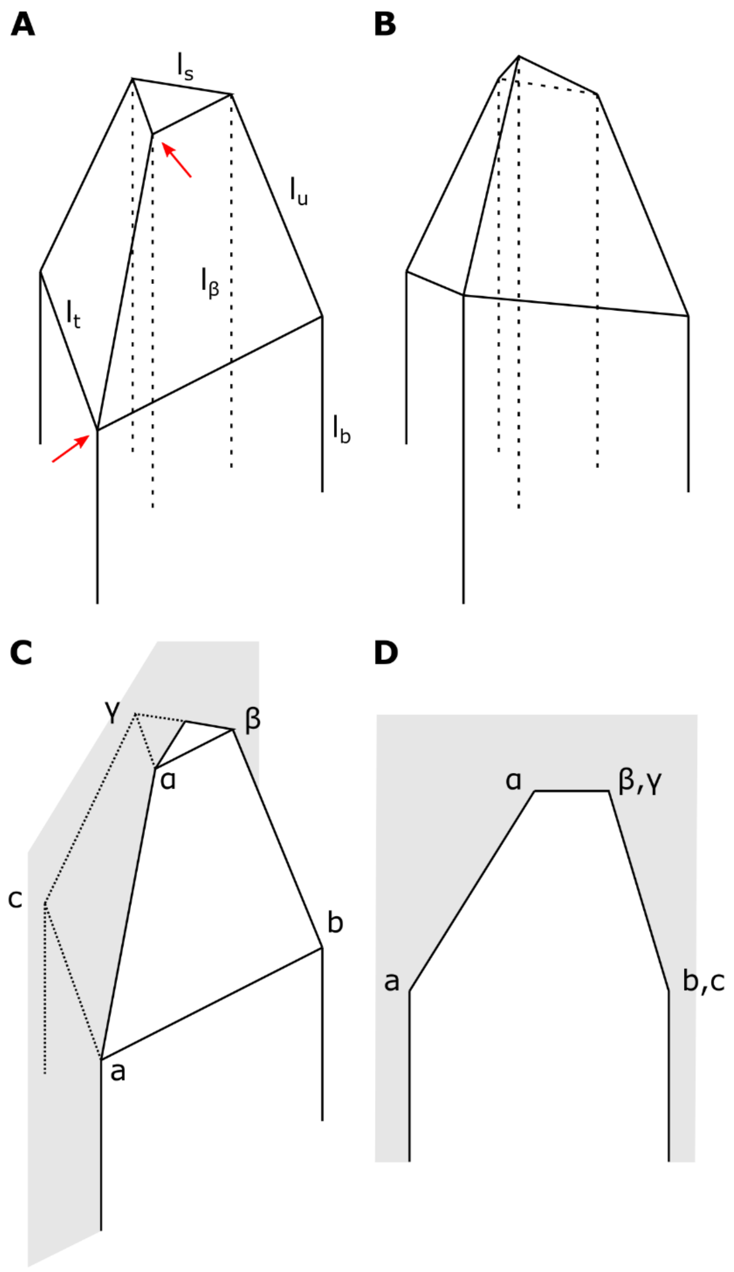

**Figure S4: Modelling representation of the ovaries**

(**A**) The ovaries are modelled as polyhedra composed of triangular prisms topped with a truncated pyramid. Red arrows denote the nodes corresponding to the top of the inner and outer walls of the expanding carpel.

(**B**) After expansion of the walls indicated by arrows in (**A**), a polyhedron appears skewed.

(**C**, **D**) Schematic of the plane over which the polyhedron is projected. The projection plane (in grey) passes through nodes *a* and *α*, and intersects the opposite edges forming a 90 degree angle (**C**). Lateral view of the projected polyhedron (**D**).

*α*, *β*, *γ* indicate the nodes at the base of the style, and *a*, *b*, *c* the nodes where the ovary roughly ceases to be vertically straight relative to the base of the ovary (**C**, **D**). $l_a$ to $l_\gamma$ indicate the shortest lengths of the segments connecting the respective nodes to the base of the ovary. Only $l_b$ and $l_\beta$ are shown for clarity.

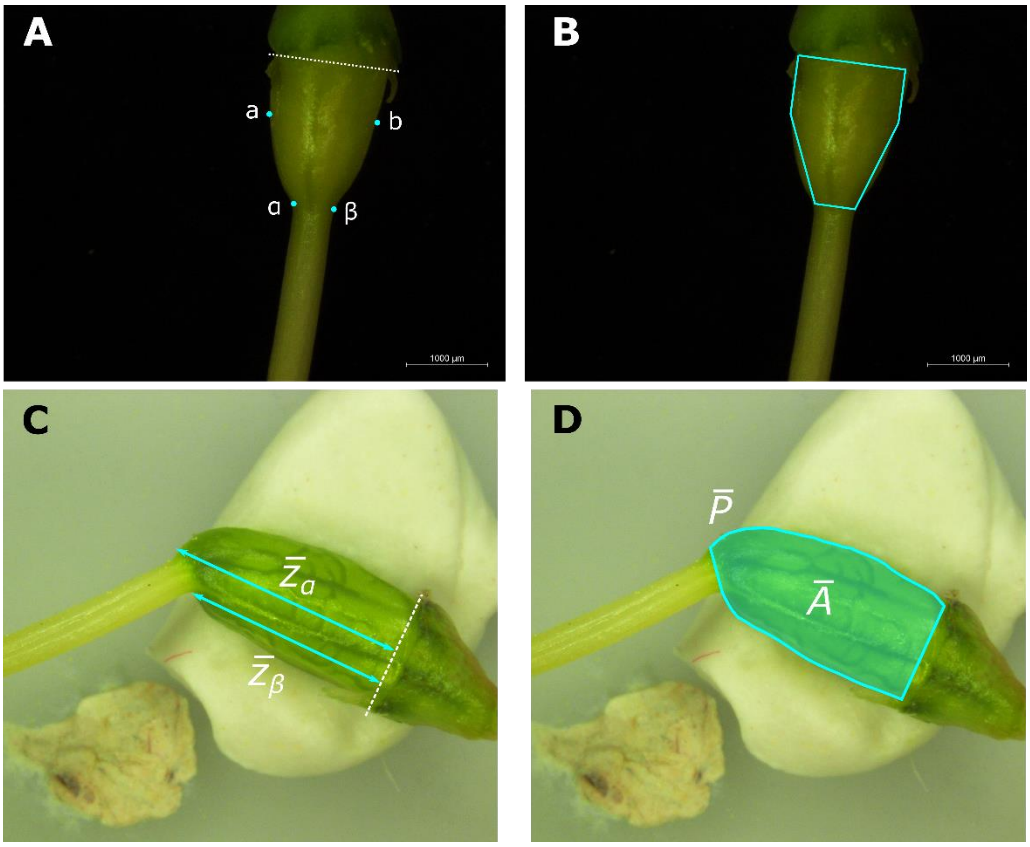

**Figure S5: Morphology of ovaries at early and late stage of development.**

(**A**, **B**) In this early-stage ovary, nodes *a* and *b* are identified as roughly the location when the ovary starts bending towards the style as observed from the microscopy image. The nodes *α* and β are at the base of the style (**A**). Based on *a*, *b*, *α* and β, the initial polyhedron is drawn (**B**).

(**C**, **D**) In this late-stage ovary, the blue lines represent the measure of the distance between the base of the ovary and the base of the style, at its extremes (**C**). The solid line represents the perimeter of the ovary, and the opaque area is used to measure the area of the ovary (**D**).

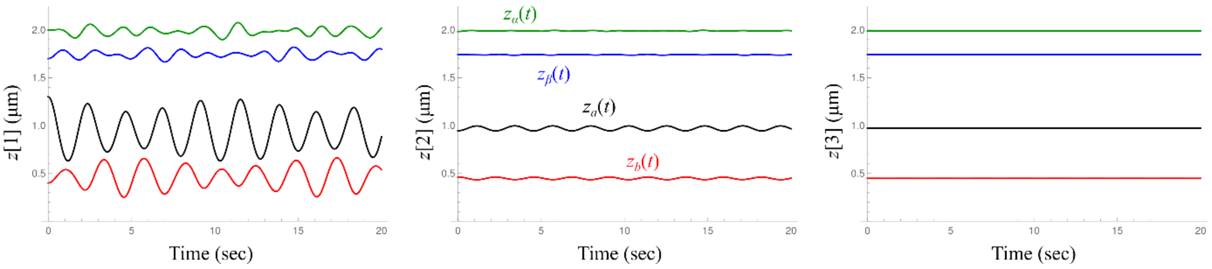

**Figure S6: Solutions of ODEs.**

Solution of the system of ODEs Eqs. (S10) after the (from left to right) first, second, and third iteration for a generic set of initial conditions. In every plot from top to bottom, we show the solutions for (green) $z_\alpha(t)$, (blue) $z_\beta(t) = z_\gamma(t)$, (black) $z_a(t)$, and (red) $z_b(t) = z_c(t)$.

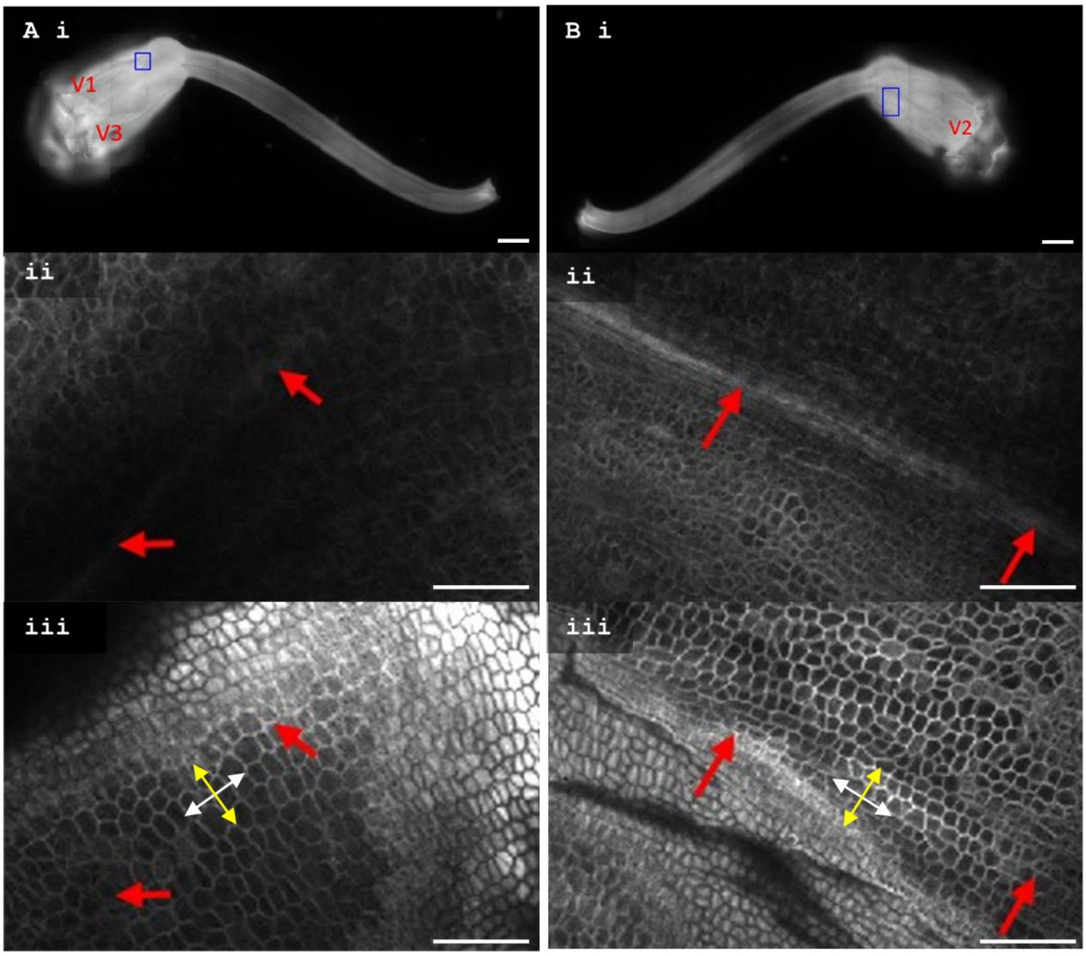

**Figure S7: Illustration of cell length and width measurements in *Cyanella alba* subsp*. flavescens***

(**A**, **B**). A left-handed *C. alba* pistil stained with Calcoflour white is imaged from both sides Overviews (panels **i**) were generated using tile scanning (20% overlap) captured on a widefield Zeiss monochrome CCD camera. From view (**A**), veins 1 and 3 are visible (marked), whereas in view (**B**) vein 2 can be seen. Blue boxes indicate the approximate position of z-stacks **ii-iii**. The midveins (panels **ii**) are clearly visible just below the surface layer (panels **iii**) and are marked in the z-stack (red arrows). A line of 5-10 cells between the arrows (panels **iii**) were measured in ImageJ to calculate an average cell size measurement along vein 1 (**A**) and vein 2 (**B**). Cell length was measured in the direction indicated by the white arrows, whereas cell width was measured in the direction indicated by the yellow arrows. Scale bars in (**i**) represent 1000 µm, whereas in (**ii**) and (**iii**) they represent 100 µm.

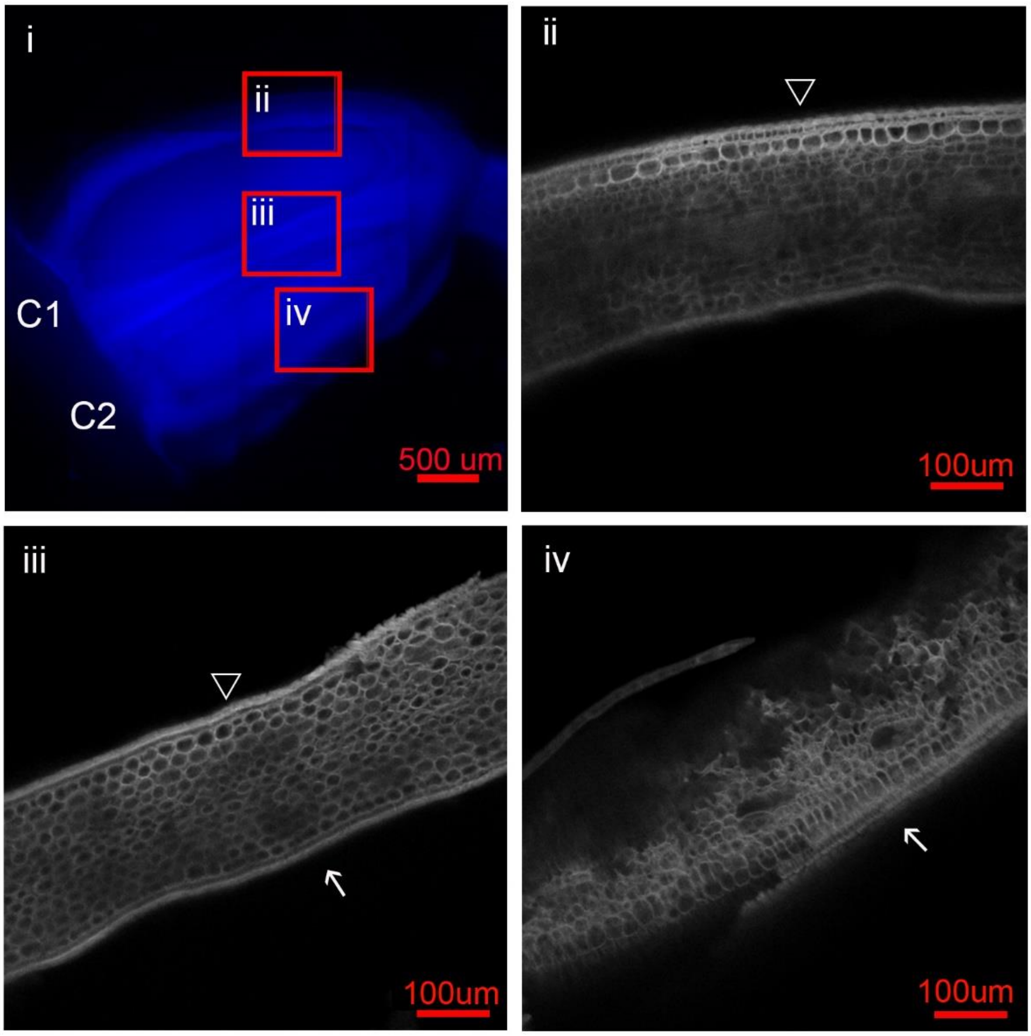

**Figure S8: Illustration of measurements of cell lengths in outer and inner carpel walls**

(**i**). A right-handed *C. alba* pistil was dissected, had its ovules removed, and stained with Calcoflour white. Red boxes indicate the approximate position of z-stacks. Selected images used for calculating cell length from these z-stacks are shown in ii-iv. A line of 10 subepidermal cells were measured in ImageJ to calculate the average cell size for the outer carpel walls (ii and iv), and inner carpel walls (iii) ∆ indicates walls of carpel 1, and the white arrow indicates the walls of carpel 2. Scale bar in (**i**) represents 500 µm, whereas in (**ii-iv**) they represent 100 µm.

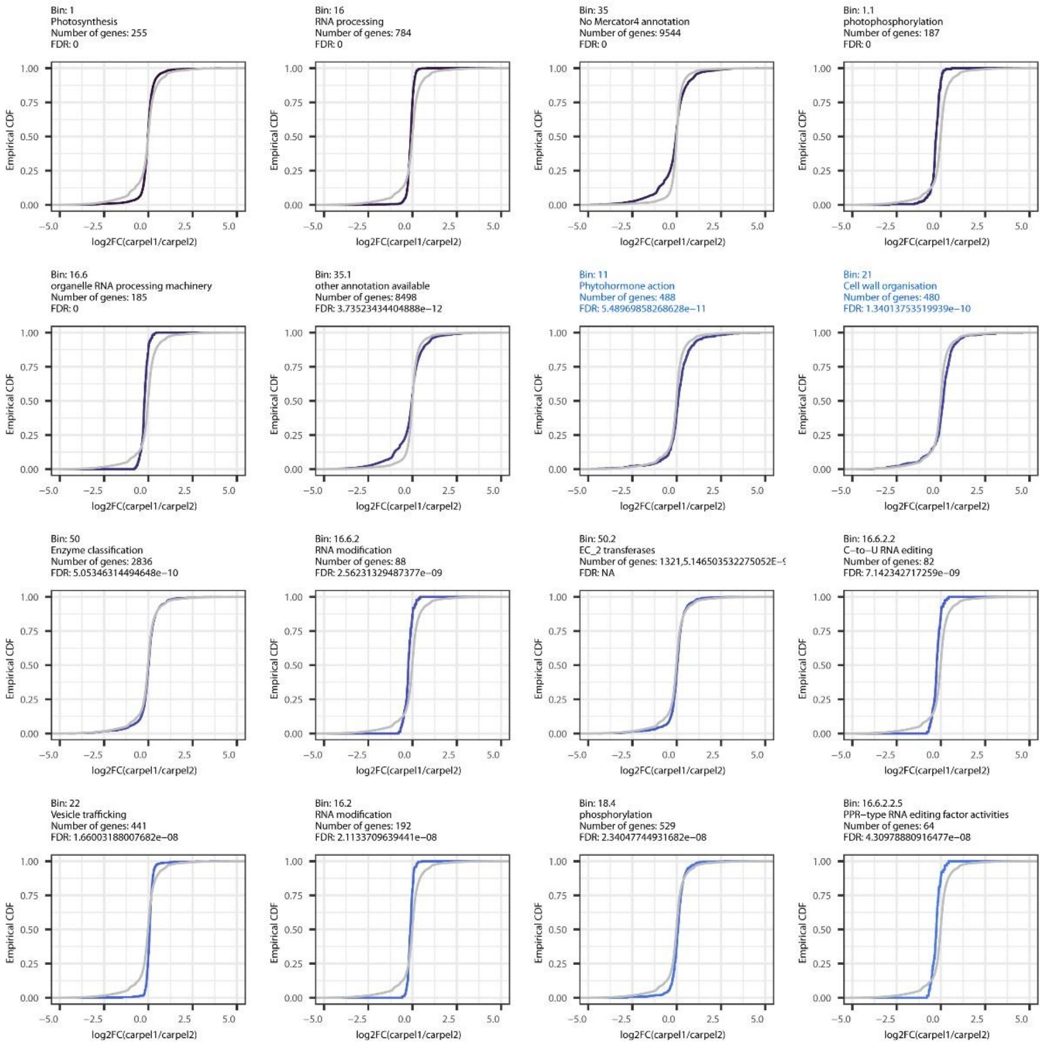
Bin: 1
Photosynthesis
Number of genes: 255
FDR: 0
Bin: 16
RNA processing
Number of genes: 784
FDR: 0
Bin: 35
No Mercator4 annotation
Number of genes: 9544
FDR: 0
Bin: 1.1
photophosphorylation
Number of genes: 187
FDR: 0
Bin: 16.6
organelle RNA processing machinery
Number of genes: 185
FDR: 0
Bin: 35.1
other annotation available
Number of genes: 8498
FDR: 3.73523434404888e−12
Bin: 11
Phytohormone action
Number of genes: 488
FDR: 5.48969858268628e−11
Bin: 21
Cell wall organisation
Number of genes: 480
FDR: 1.34013753519939e−10
Bin: 50
Enzyme classification
Number of genes: 2836
FDR: 5.05346314494648e−10
Bin: 16.6.2
RNA modification
Number of genes: 88
FDR: 2.56231329487377e−09
Bin: 50.2
EC_2 transferases
Number of genes: 1321,5.146503532275052E-[illegible]
FDR: NA
Bin: 16.6.2.2
C−to−U RNA editing
Number of genes: 82
FDR: 7.142342717259e−09
Bin: 22
Vesicle trafficking
Number of genes: 441
FDR: 1.66003188007682e−08
Bin: 16.2
RNA modification
Number of genes: 192
FDR: 2.1133709639441e−08
Bin: 18.4
phosphorylation
Number of genes: 529
FDR: 2.34047744931682e−08
Bin: 16.6.2.2.5
PPR−type RNA editing factor activities
Number of genes: 64
FDR: 4.30978880916477e−08
Empirical CDF
log2FC(carpel1/carpel2)

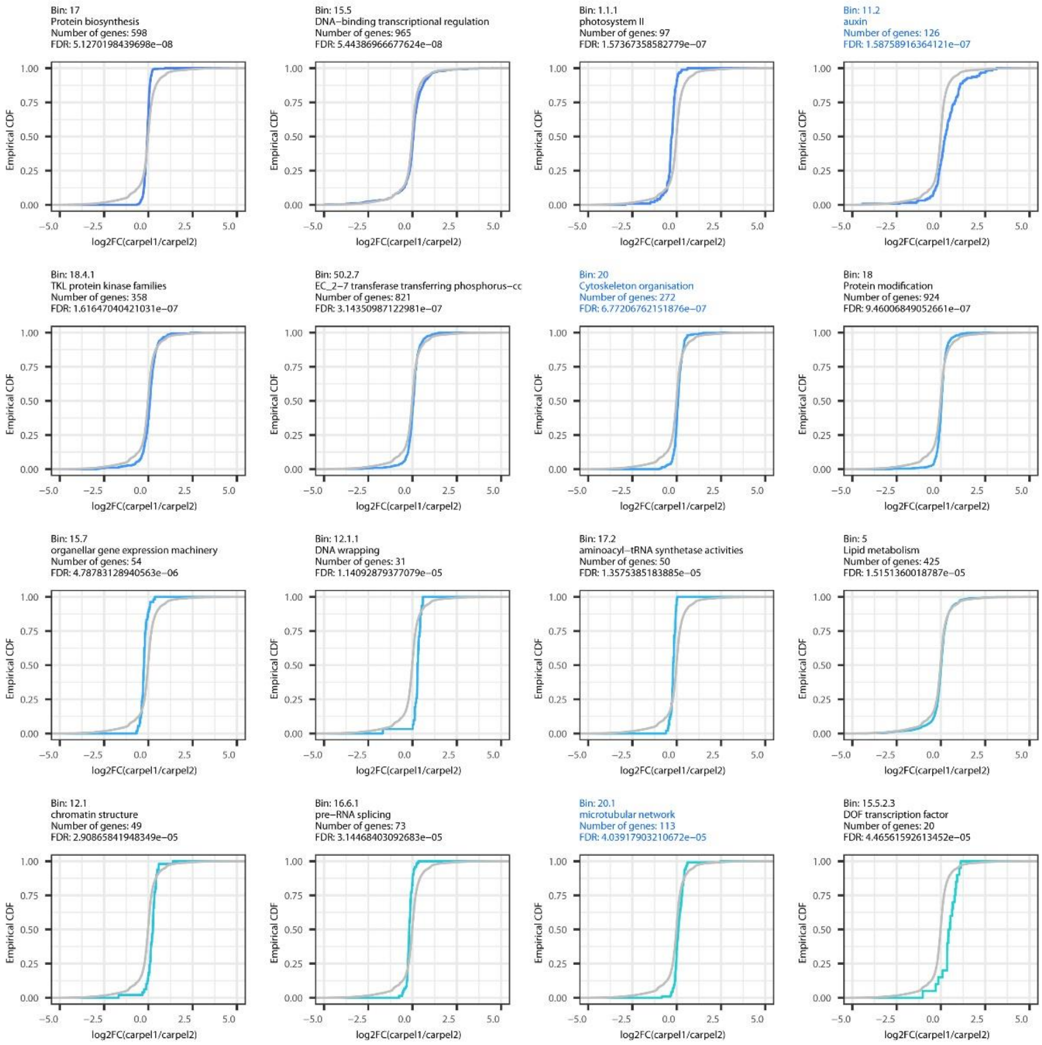

Bin: 17
Protein biosynthesis
Number of genes: 598
FDR: 5.1270198439698e−08
Bin: 15.5
DNA−binding transcriptional regulation
Number of genes: 965
FDR: 5.44386966677624e−08
Bin: 1.1.1
photosystem II
Number of genes: 97
FDR: 1.57367358582779e−07
Bin: 11.2
auxin
Number of genes: 126
FDR: 1.58758916364121e−07
Bin: 18.4.1
TKL protein kinase families
Number of genes: 358
FDR: 1.61647040421031e−07
Bin: 50.2.7
EC_2−7 transferase transferring phosphorus−cc
Number of genes: 821
FDR: 3.14350987122981e−07
Bin: 20
Cytoskeleton organisation
Number of genes: 272
FDR: 6.77206762151876e−07
Bin: 18
Protein modification
Number of genes: 924
FDR: 9.46006849052661e−07
Bin: 15.7
organellar gene expression machinery
Number of genes: 54
FDR: 4.78783128940563e−06
Bin: 12.1.1
DNA wrapping
Number of genes: 31
FDR: 1.14092879377079e−05
Bin: 17.2
aminoacyl−tRNA synthetase activities
Number of genes: 50
FDR: 1.3575385183885e−05
Bin: 5
Lipid metabolism
Number of genes: 425
FDR: 1.5151360018787e−05
Bin: 12.1
chromatin structure
Number of genes: 49
FDR: 2.90865841948349e−05
Bin: 16.6.1
pre−RNA splicing
Number of genes: 73
FDR: 3.14468403092683e−05
Bin: 20.1
microtubular network
Number of genes: 113
FDR: 4.03917903210672e−05
Bin: 15.5.2.3
DOF transcription factor
Number of genes: 20
FDR: 4.46561592613452e−05
Empirical CDF
log2FC(carpel1/carpel2)

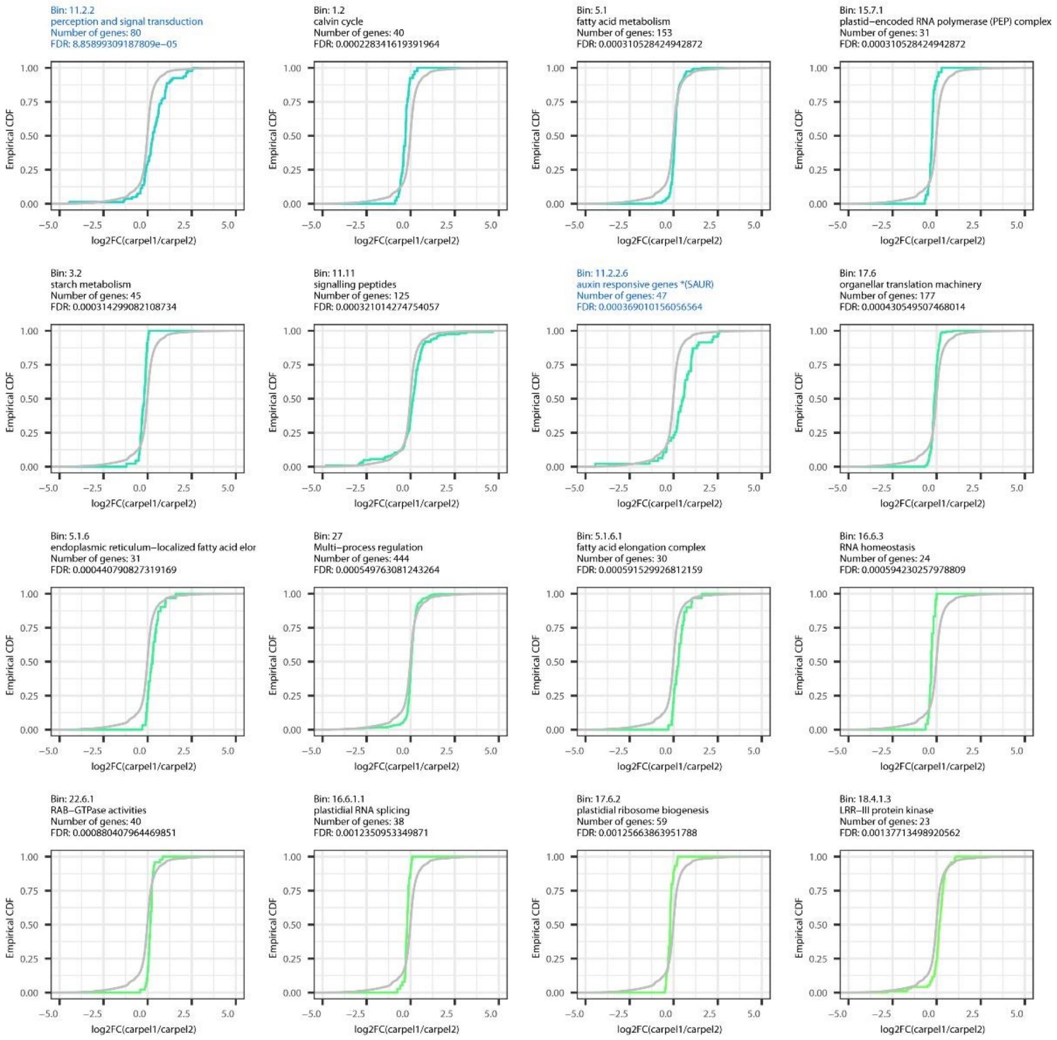
Bin: 11.2.2
perception and signal transduction
Number of genes: 80
FDR: 8.85899309187809e−05
Bin: 1.2
calvin cycle
Number of genes: 40
FDR: 0.000228341619391964
Bin: 5.1
fatty acid metabolism
Number of genes: 153
FDR: 0.000310528424942872
Bin: 15.7.1
plastid−encoded RNA polymerase (PEP) complex
Number of genes: 31
FDR: 0.000310528424942872
Bin: 3.2
starch metabolism
Number of genes: 45
FDR: 0.000314299082108734
Bin: 11.11
signalling peptides
Number of genes: 125
FDR: 0.000321014274754057
Bin: 11.2.2.6
auxin responsive genes *(SAUR)
Number of genes: 47
FDR: 0.000369010156056564
Bin: 17.6
organellar translation machinery
Number of genes: 177
FDR: 0.000430549507468014
Bin: 5.1.6
endoplasmic reticulum−localized fatty acid elor
Number of genes: 31
FDR: 0.000440790827319169
Bin: 27
Multi−process regulation
Number of genes: 444
FDR: 0.000549763081243264
Bin: 5.1.6.1
fatty acid elongation complex
Number of genes: 30
FDR: 0.000591529926812159
Bin: 16.6.3
RNA homeostasis
Number of genes: 24
FDR: 0.000594230257978809
Bin: 22.6.1
RAB−GTPase activities
Number of genes: 40
FDR: 0.000880407964469851
Bin: 16.6.1.1
plastidial RNA splicing
Number of genes: 38
FDR: 0.0012350953349871
Bin: 17.6.2
plastidial ribosome biogenesis
Number of genes: 59
FDR: 0.00125663863951788
Bin: 18.4.1.3
LRR−III protein kinase
Number of genes: 23
FDR: 0.00137713498920562
Empirical CDF
log2FC(carpel1/carpel2)

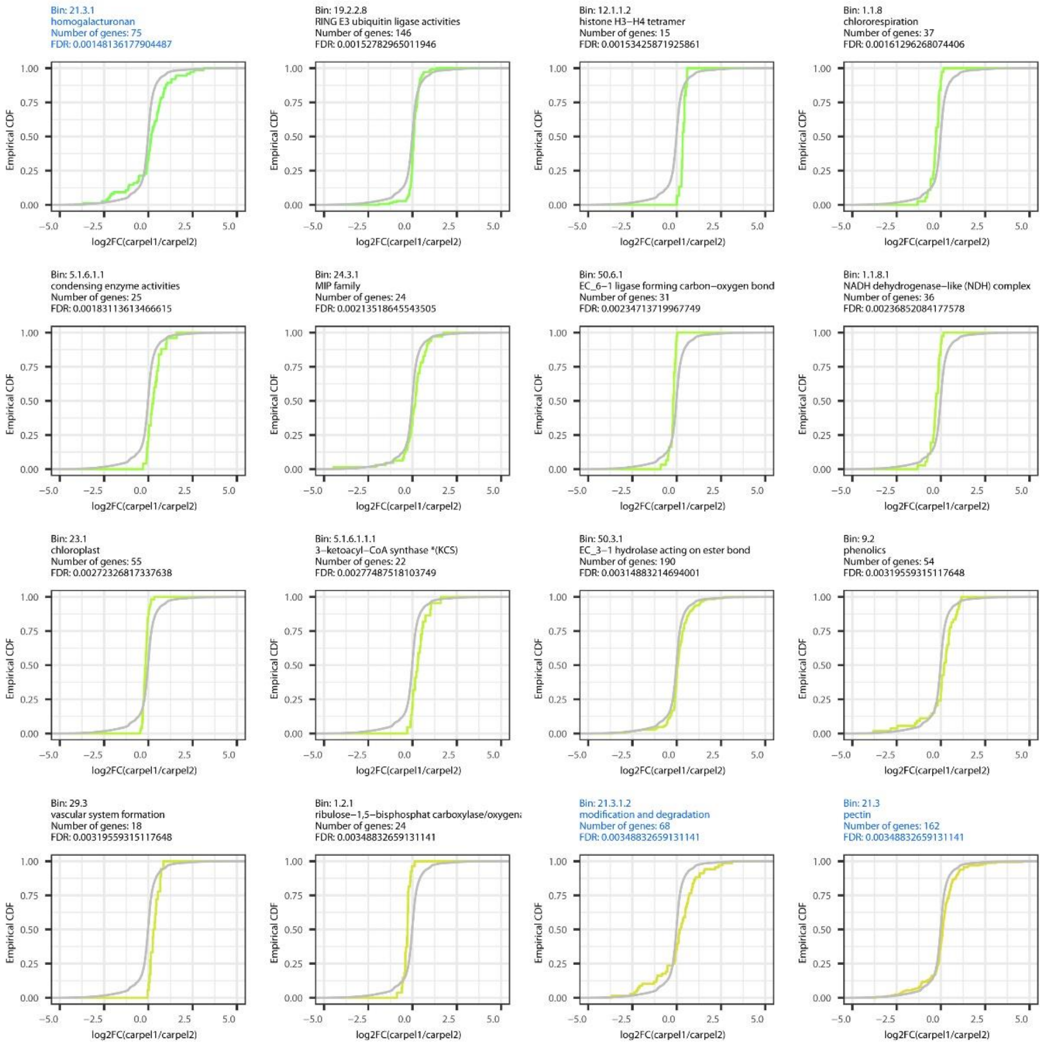

Bin: 21.3.1
homogalacturonan
Number of genes: 75
FDR: 0.00148136177904487
Bin: 19.2.2.8
RING E3 ubiquitin ligase activities
Number of genes: 146
FDR: 0.00152782965011946
Bin: 12.1.1.2
histone H3–H4 tetramer
Number of genes: 15
FDR: 0.00153425871925861
Bin: 1.1.8
chlororespiration
Number of genes: 37
FDR: 0.00161296268074406
Bin: 5.1.6.1.1
condensing enzyme activities
Number of genes: 25
FDR: 0.00183113613466615
Bin: 24.3.1
MIP family
Number of genes: 24
FDR: 0.00213518645543505
Bin: 50.6.1
EC_6–1 ligase forming carbon–oxygen bond
Number of genes: 31
FDR: 0.00234713719967749
Bin: 1.1.8.1
NADH dehydrogenase–like (NDH) complex
Number of genes: 36
FDR: 0.00236852084177578
Bin: 23.1
chloroplast
Number of genes: 55
FDR: 0.00272326817337638
Bin: 5.1.6.1.1.1
3–ketoacyl–CoA synthase *(KCS)
Number of genes: 22
FDR: 0.00277487518103749
Bin: 50.3.1
EC_3–1 hydrolase acting on ester bond
Number of genes: 190
FDR: 0.00314883214694001
Bin: 9.2
phenolics
Number of genes: 54
FDR: 0.00319559315117648
Bin: 29.3
vascular system formation
Number of genes: 18
FDR: 0.00319559315117648
Bin: 1.2.1
ribulose–1,5–bisphosphat carboxylase/oxygena
Number of genes: 24
FDR: 0.00348832659131141
Bin: 21.3.1.2
modification and degradation
Number of genes: 68
FDR: 0.00348832659131141
Bin: 21.3
pectin
Number of genes: 162
FDR: 0.00348832659131141
Empirical CDF
log2FC(carpel1/carpel2)

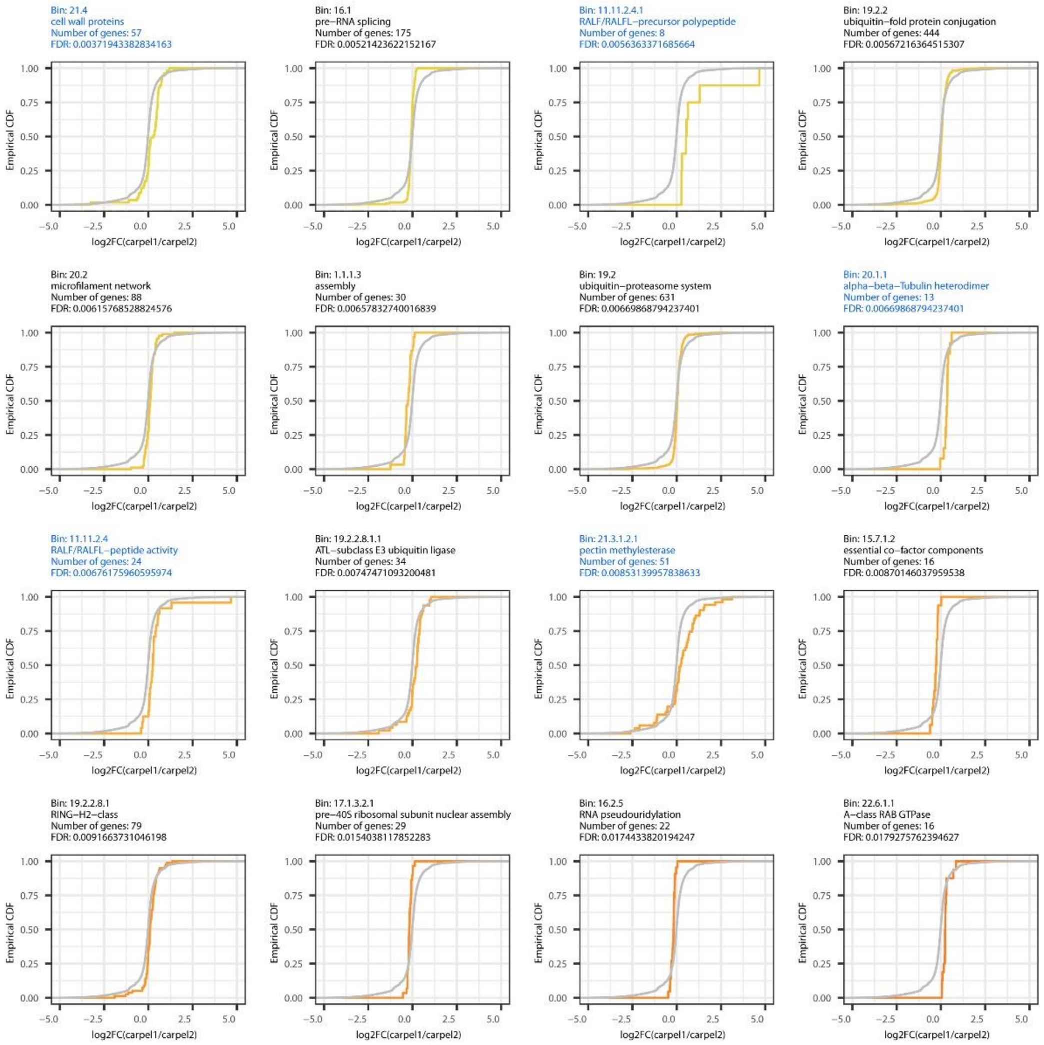
Bin: 21.4
cell wall proteins
Number of genes: 57
FDR: 0.00371943382834163
Bin: 16.1
pre–RNA splicing
Number of genes: 175
FDR: 0.00521423622152167
Bin: 11.11.2.4.1
RALF/RALFL–precursor polypeptide
Number of genes: 8
FDR: 0.0056363371685664
Bin: 19.2.2
ubiquitin–fold protein conjugation
Number of genes: 444
FDR: 0.00567216364515307
Bin: 20.2
microfilament network
Number of genes: 88
FDR: 0.00615768528824576
Bin: 1.1.1.3
assembly
Number of genes: 30
FDR: 0.00657832740016839
Bin: 19.2
ubiquitin–proteasome system
Number of genes: 631
FDR: 0.00669868794237401
Bin: 20.1.1
alpha–beta–Tubulin heterodimer
Number of genes: 13
FDR: 0.00669868794237401
Bin: 11.11.2.4
RALF/RALFL–peptide activity
Number of genes: 24
FDR: 0.00676175960595974
Bin: 19.2.2.8.1.1
ATL–subclass E3 ubiquitin ligase
Number of genes: 34
FDR: 0.00747471093200481
Bin: 21.3.1.2.1
pectin methylesterase
Number of genes: 51
FDR: 0.00853139957838633
Bin: 15.7.1.2
essential co–factor components
Number of genes: 16
FDR: 0.00870146037959538
Bin: 19.2.2.8.1
RING–H2–class
Number of genes: 79
FDR: 0.0091663731046198
Bin: 17.1.3.2.1
pre–40S ribosomal subunit nuclear assembly
Number of genes: 29
FDR: 0.0154038117852283
Bin: 16.2.5
RNA pseudouridylation
Number of genes: 22
FDR: 0.0174433820194247
Bin: 22.6.1.1
A–class RAB GTPase
Number of genes: 16
FDR: 0.0179275762394627
Empirical CDF
log2FC(carpel1/carpel2)

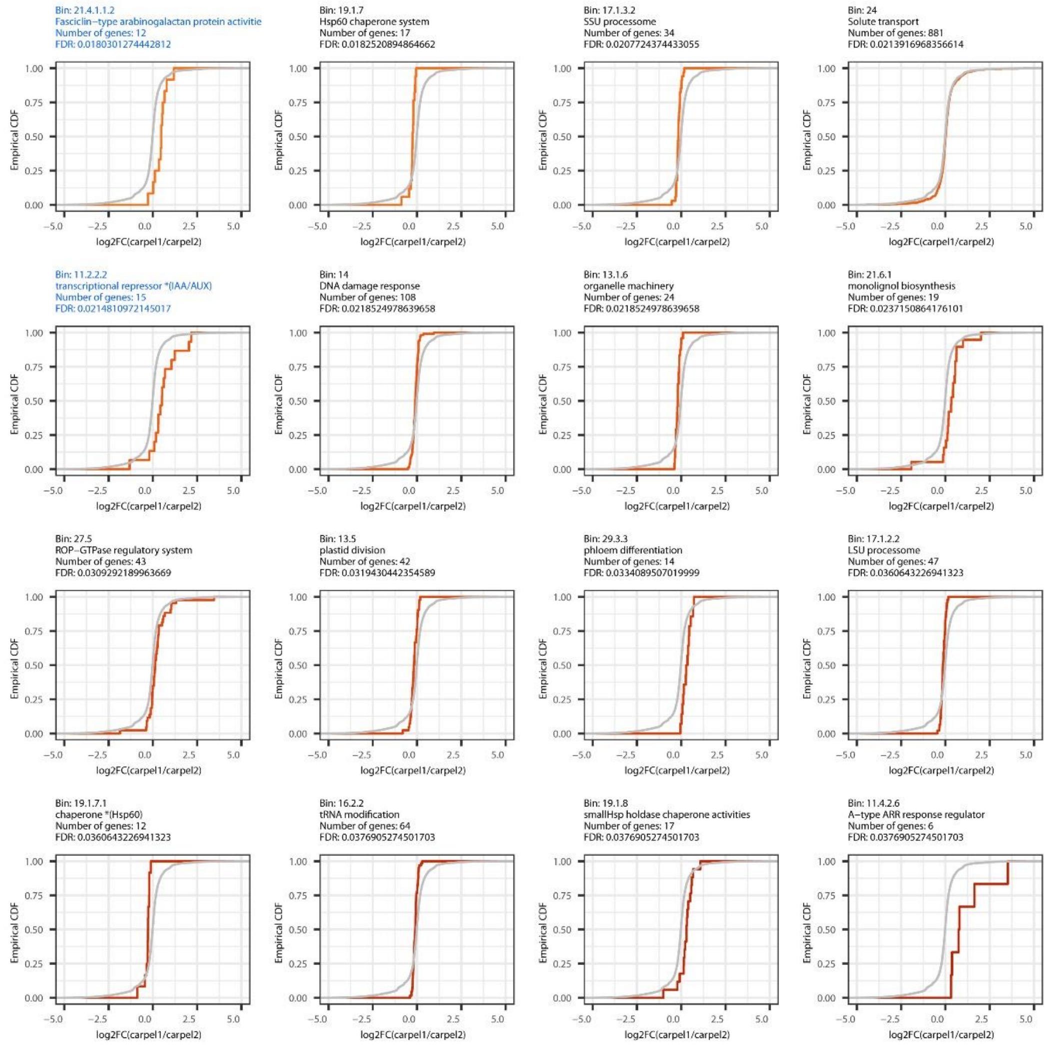
Bin: 21.4.1.1.2
Fasciclin–type arabinogalactan protein activitie
Number of genes: 12
FDR: 0.0180301274442812
Bin: 19.1.7
Hsp60 chaperone system
Number of genes: 17
FDR: 0.0182520894864662
Bin: 17.1.3.2
SSU processome
Number of genes: 34
FDR: 0.0207724374433055
Bin: 24
Solute transport
Number of genes: 881
FDR: 0.0213916968356614
Bin: 11.2.2.2
transcriptional repressor *(IAA/AUX)
Number of genes: 15
FDR: 0.0214810972145017
Bin: 14
DNA damage response
Number of genes: 108
FDR: 0.0218524978639658
Bin: 13.1.6
organelle machinery
Number of genes: 24
FDR: 0.0218524978639658
Bin: 21.6.1
monolignol biosynthesis
Number of genes: 19
FDR: 0.0237150864176101
Bin: 27.5
ROP–GTPase regulatory system
Number of genes: 43
FDR: 0.0309292189963669
Bin: 13.5
plastid division
Number of genes: 42
FDR: 0.0319430442354589
Bin: 29.3.3
phloem differentiation
Number of genes: 14
FDR: 0.0334089507019999
Bin: 17.1.2.2
LSU processome
Number of genes: 47
FDR: 0.0360643226941323
Bin: 19.1.7.1
chaperone *(Hsp60)
Number of genes: 12
FDR: 0.0360643226941323
Bin: 16.2.2
tRNA modification
Number of genes: 64
FDR: 0.0376905274501703
Bin: 19.1.8
smallHsp holdase chaperone activities
Number of genes: 17
FDR: 0.0376905274501703
Bin: 11.4.2.6
A–type ARR response regulator
Number of genes: 6
FDR: 0.0376905274501703
Empirical CDF
log2FC(carpel1/carpel2)

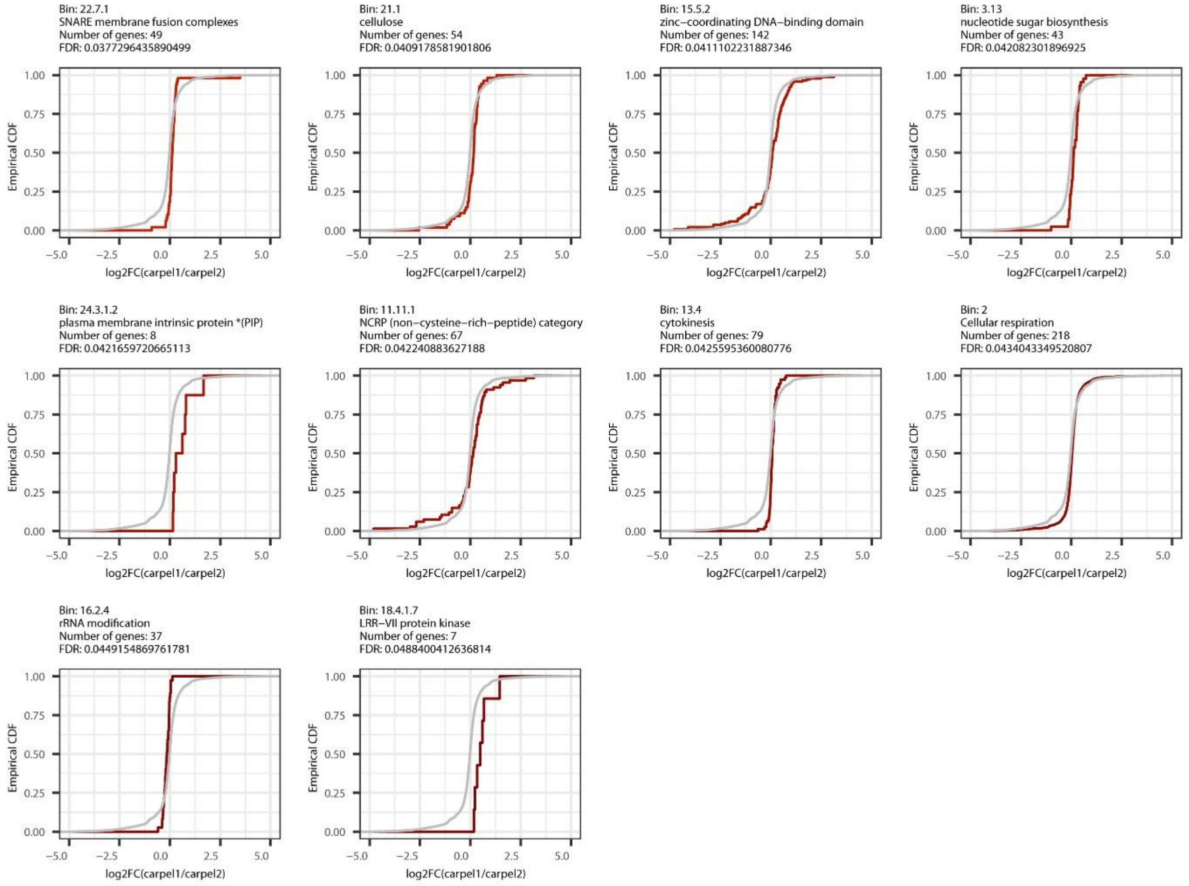
Bin: 22.7.1
SNARE membrane fusion complexes
Number of genes: 49
FDR: 0.0377296435890499
Bin: 21.1
cellulose
Number of genes: 54
FDR: 0.0409178581901806
Bin: 15.5.2
zinc–coordinating DNA–binding domain
Number of genes: 142
FDR: 0.0411102231887346
Bin: 3.13
nucleotide sugar biosynthesis
Number of genes: 43
FDR: 0.042082301896925
Bin: 24.3.1.2
plasma membrane intrinsic protein *(PIP)
Number of genes: 8
FDR: 0.0421659720665113
Bin: 11.11.1
NCRP (non–cysteine–rich–peptide) category
Number of genes: 67
FDR: 0.042240883627188
Bin: 13.4
cytokinesis
Number of genes: 79
FDR: 0.0425595360080776
Bin: 2
Cellular respiration
Number of genes: 218
FDR: 0.0434043349520807
Bin: 16.2.4
rRNA modification
Number of genes: 37
FDR: 0.0449154869761781
Bin: 18.4.1.7
LRR–VII protein kinase
Number of genes: 7
FDR: 0.0488400412636814
Empirical CDF
log2FC(carpel1/carpel2)

**Figure S9: Mercator bins identified as having significantly different distributions of log2 fold-changes in expression between carpels 1 and 2 of *Cyanella alba* subsp*. flavescens***

ECDF plots are shown for all bins that were identified as significantly different from the transcriptomic background. For each bin, the bin number, short description, number of genes contained in the bin and FDR value is indicated, and the grey line is the transcriptomic background (all transcripts outside of the bin). Bins discussed in the main text are highlighted with blue font.

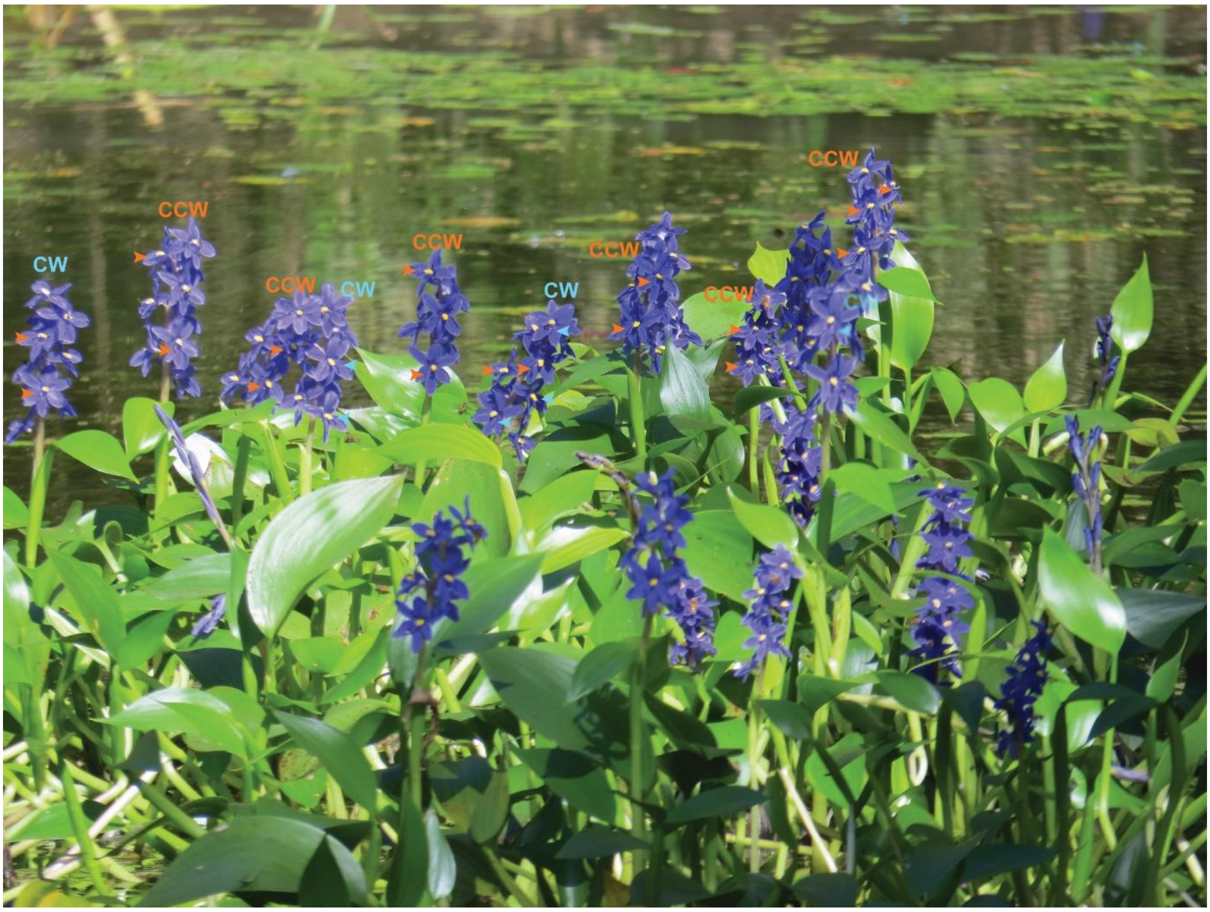

**Figure S10: *Monochoria australasica* inflorescences, a species with inflorescence-level monomorphic enantiostyly**

Phyllotaxis orientation is indicated as CW (clockwise) or CCW (counter clockwise). Orange and blue arrowheads point to styles deflected to the left (orange) or right (blue). See Table S11.

### Supporting Information Tables

**Table S1: Numbers of 10-kb windows in the genome with different coverage between L- and R-plants in Illumina2021 and Illumina2022 datasets**

See Supplemental Excel file with Tables S1, S2, S9, S10.

**Table S2: Differentially expressed transcripts between styles of L- and R-plants at each developmental stage**

See Supplemental Excel file with Tables S1, S2, S9, S10.

**Tables S3 to S8:** See section on biomechanical model above in Supporting Information Text

**Table S9: Differentially expressed genes between carpels 1 and 2**

See Supplemental Excel file with Tables S1, S2, S9, S10.

**Table S10: Mercator bins with significant difference in carpel 1/carpel 2 expression compared to genomic background**

See Supplemental Excel file with Tables S1, S2, S9, S10.

**Table S11: List of available *Monochoria austalasica* images with their phyllotaxis orientation and style deflection.**

CW: clockwise, CCW: counterclockwise, when moving from older to younger organs.

| Image link | Phyllotaxis | Style deflection |
|---|---|---|
| https://www.inaturalist.org/observations/74107844 | CCW | L |
| https://www.inaturalist.org/observations/65301963 | CW | R |
| https://www.inaturalist.org/observations/60589407 | CCW | L |
| https://www.inaturalist.org/observations/40130455 | CW | R |
| https://www.inaturalist.org/observations/19364608 | CW | R |
| | CW | R |
| | CW | R |
| https://www.inaturalist.org/observations/19364608 (large image; see Figure S10) | CW | L |
| | CCW | L |
| | CCW | L |
| | CW | R |
| | CCW | L |
| | CW | mixed |
| | CCW | L |
| | CCW | L |
| | CCW | L |
| | CW | R |
| SCHB image | CCW | L |
| https://www.flickr.com/photos/71736071@N02/34452474370/ | CCW | L |

| | | |
|---|---|---|
| https://www.inaturalist.org/observations/151569011 (misidentified as *P. cyanea*) | CCW | L |
| https://www.inaturalist.org/observations/121835877 (misidentified as *P. cyanea*) | CCW | L |

### Supporting Information Materials and Methods

#### Plant study system and sites

*Cyanella alba* subsp. *flavescens* is a narrow endemic native to the Western Cape of South Africa. It is restricted to the northern Cederberg and Olifants River region and is locally abundant in the Biedouw River Valley and at Wupperthal (see Fig 15 in Manning and Goldblatt, 2012). The species is a long-lived, deciduous geophyte with a deep-seated corm and pale-yellow enantiostylous zygomorphic flowers with styles deflected to the left and right side of the flower (see Figure 1A in main text). Anthers are dimorphic (heteranthery) with poricidal dehiscence. The upper five centrally placed “feeding anthers” are fused and the sixth “pollinating anther” is deflected downwards to either the left or right side of the flower in an opposite direction to the style.

Populations in the Biedouw River Valley and Wupperthal are large (estimated at 2000-5000 individuals) and flower from early August to October after winter rains. The three populations we sampled for demographic studies (see below) in the 2022 and 2023 flowering seasons were located in the Biedouw Valley (site 1 at 32°08'18" E19°10'57", 400 m a.s.l and site 2 at S32°11'25" E19°10'17", 560 m a.s.l.) and Wupperthal (site 3 at S32°16'17" E19°12'42", 530 m a.s.l.). All sites occurred on open degraded pasture comprised of renosterveld vegetation and all sites were separated by a minimum of ~10km. Our research was approved by Cape Nature (Permit CN35-87-25844) and we collected plants with permission from Mr Barry Lubbe of Mertenhof farm and Barend Salomo of the Wupperthal Original Rooibos Cooperative.

#### Demographic study.

To investigate the dynamics of floral handedness in *C. alba flavescens* throughout two flowering seasons, we marked 75 plants at each of the three sites with a unique number at the beginning of 2022. The number was etched into a metal tag to withstand weathering and secured next to the plant using a 10 cm nail driven into the ground. We recorded the handedness of flowers on each plant across all three sites every one to two days during the 2022 flowering season, and every four to five days during 2023. At each visit, newly opened flowers were recorded as L or R and marked individually with a jeweller’s tag. Flowers of *C. alba flavescens* remain open on average for 8.9 days, range 5-12 days (B. Paudel et al., unpubl. data, 2024), In 2023 we modified our methods for plants with more than one flower open by recording the position of the flowers within the inflorescence and assigning those closest to the ground as having opened first. This more accurately estimated the sequence of handedness in the inflorescence. We also recorded for all plants the number of buds, open and wilted flowers at each visit.

To identify the hypothesized *E*-locus controlling L- or R- handedness in 2022 we collected two to three leaves from 16 individuals from sites 1 (*n*=8), 2 (*n*=5), and 3 (*n*=3) that produced six flowers that were either R or L only. The leaves were dried in silica before gDNA extraction and sequencing as described below.

**DNA extraction and sequencing.**

We extracted total genomic DNA from silica dried or frozen leaf material using Nucleobond High Molecular Weight DNA extraction kits (Macherey-Nagel™, cat no.: 740_160.20) and samples were sent to Novogene UK (Cambridge Sequencing Centre) for whole genome sequencing on the Illumina platform (NovaSeq 6000). A single, right-handed sample (C_209) was submitted to Novogene UK for high fidelity PacBio sequencing (PacBio sequel II/IIe DNA HiFi library).

**RNA extraction and sequencing.**

We extracted RNA from *C. alba flavescens* pistils, including the style and half of the ovary, at three developmental stages: “mid”, “early” and “very early” using a basic phenol:chloroform extraction method followed by purification with an RNA Miniprep kit (Zymo Research: Direct-zol RNA Miniprep Kits; cat. no. R2050). We assigned buds to bud classes based on a number of visual criteria detailed in Figure 3A, and all open flowers and dissected buds on an individual plant were examined before assigning a handedness phenotype (L or R) to a sample. We pooled dissected pistils at each developmental stage across multiple individuals to generate three biological replicates. Mid and early-stage samples consisted of five to seven pistils (~30 mg tissue) and very early samples consisted of ten pistils (~20 mg tissue). RNA was sequenced on the Illumina platform following Eukaryotic mRNA library construction (polyA enrichment).

We also extracted RNA from the outer walls of carpel 1 and carpel 2 of *C. alba*. We divided the individuals into four groups and pooled the dissected outer walls of carpel 1 and carpel 2 in each group separately to generate four pairs of biological replicates. Each sample consisted of ~15 carpels. The RNA was extracted and sequenced using the same methods described in the preceding paragraph.

**PacBio Assembly.**

We used the *k*-mer counting tool KMC version 3.2.1 (Kokot et al., 2017) to count *21*-mers in the PacBio sequences of the sample C_209. Then we used GenomeScope2.0 (Ranallo-Benavidez et al., 2020) to estimate the genome size and ploidy of *C. alba flavescens*.

The assembly was generated using hifiasm version 0.19.0-r534 (Cheng et al., 2021) and we used default settings with only “--hg-size” (estimated haploid genome size) specified. Haploid genome size estimates of 1G and 700M were used, in line with flow cytometry and GenomeScope2.0 size estimates, and we estimated completeness and contiguity of the assemblies using BUSCO version 5.4.4 and QUAST version 5.2.0 (Manni et al., 2021; Mikheenko et al., 2018). The database chosen for the BUSCO analysis was liliopsida_odb10. The “-m genome” and “--augustus” options were specified.

We downloaded the *Arabidopsis* protein database "Araport11_pep_20220914" from the website ‘The *Arabidopsis* Information Resource’ (TAIR, www.arabidopsis.org). We used tblastn to align the *Arabidopsis* protein sequences to the reference genome, to identify protein-coding regions in the genome. Bedtools version 2.30.0 (Quinlan and Hall, 2010) was used to merge identified regions that are less than 10 bp apart.

**Genome Wide Association Study (GWAS).**

We used NextGenMap version 0.5.5 to align Illumina reads to the primary PacBio assembly (Sedlazeck et al., 2013). We used SAMtools version 1.3.1 (Danecek et al., 2021) to exclude reads with mapping quality lower than 10, then sort and index the alignment files. We used the function “mpileup” of BCFtools version 1.16 (Danecek et al., 2021) to generate binary variant call files (BCFs) with the alignment files. The “-Q 10” options were specified to skip bases with base quality lower than 10, and only protein-coding regions in the reference genome were included. We used the function “call” of BCFtools to identify potential variant sites with the multiallelic calling model option (-m) included. We used VCFtools version 0.1.16 (Danecek et al., 2011) to label genotypes with coverage < 3 as “missing”, and filter sites with minor allele count lower than 3, mapping quality < 30, or more than 50% “missing” genotypes (options: --minDP 3; --mac 3; --minQ 30; --max-missing 0.5). We used BCFtools to keep only single-nucleotide polymorphisms (SNPs) (option: --types snps). Finally, we used PLINK version 1.9.0 (http://pngu.mgh.harvard.edu/purcell/plink/) to identify SNPs significantly associated with L or R morphs (Cochran-Armitage test, $P<0.0005$).

**Coverage analysis.**

We performed coverage analyses with non-overlapping genomic windows of 10 kb. Illumina reads were aligned to the two haplotypes of the genome assembly using NextGenMap and analysed separately. SAMtools was used to remove reads with a mapping quality less than 10 before sorting and indexing the alignment files. We calculated read coverage over 10-kb windows using mosdepth version 0.3.3 (Pedersen and Quinlan, 2018) and coverage of each window was normalised by dividing by the genome-wide average read coverage. We analysed normalised coverage scores in R version 4.3.0 and *t*-tests were performed to determine whether the normalised coverage scores of left- and right-handed samples were significantly different for each window, with a cut-off of $P<0.0005$. The ratio of normalised coverage for L versus R samples was calculated, and filtered for windows where this ratio was > 1.5 or < 0.67, indicating lower coverage in one morph compared to the other. We used IGV version 2.14.1 (Thorvaldsdottir et al., 2013) to investigate regions of interest.

**KmerGO2.**

We used KmerGO2 (Wang et al., 2020, https://github.com/ChnMasterOG/KmerGO2) to compare the *k*-mer composition of L- and R-individuals to identify *k*-mers characteristic of each morph. We used *k*=40 and ran subcommands "kmc3" and "filtering" with 24 threads and "union" with 8 threads.

We imported the output of KmerGO2 into Microsoft Excel for further analysis and calculated the total number of morph-associated *k*-mers (Average of Sensitivity and Specificity (ASS) score greater than 0.8) for each dataset. We combined presence/absence scores for each *k*-mer for the forward and reverse fasta file of each individual and the total number of L-associated and R-associated *k*-mers was plotted in Excel.

**Transcriptomic analysis.**

We assembled a draft transcriptome using the combined reads from the 18 RNA samples of pistils. The *de novo* assembly was created using Trinity version 2.14.0 (Grabherr et al., 2011). The "--max_memory" argument was set to 50G and the "normalise_by_read_set" parameter included to limit RAM usage. We conducted functional annotation for the draft transcriptome using Mercator4 v6.0 (Schwacke et al., 2019) with Prot-scriber and Swissprot annotations included. We used the read aligner Salmon (version 1.9.0; (Patro et al., 2017)) with the "--no_bowtie" option and we performed read pseudoalignment and quantification with Salmon. We used the Salmon "quant" command with the "--gcBias" and "--validateMappings" settings. Count data was analysed in R-studio (R version 4.3.0) using the package DESeq2 (Love et al., 2014). Graphs were plotted using the R package ggplot2 (Wickham, 2009). For each gene, the functional annotation of the longest isoform was selected as the annotation of the gene.

We aligned the RNA-Seq data from 18 pistil samples and eight carpel samples to the reference genome with STAR (Dobin et al., 2013). Then we conducted the structural annotation of the reference genome using BRAKER3 (Gabriel et al., 2023), with the aligned RNA reads and the partition Viridiplantae of the protein database OrthoDB v11 (Kuznetsov et al., 2023) as evidence. We conducted functional annotation for the coding sequences predicted by BRAKER3 using Mercator4 v6.0 (Schwacke et al., 2019) with Prot-scriber and Swissprot annotations included. Since *SMALL AUXIN UP-REGULATED* (SAUR) genes were not included in the Mercator bins, we manually created a Mercator bin: 11.2.2.6, “Phytohormone action.auxin.perception and signal transduction.auxin responsive genes *(SAUR)”, for genes that were annotated as SAUR genes.

We quantified the expression level of each predicted gene in the eight carpel samples with StringTie (Pertea et al., 2015). We conducted gene differential expression analysis with the count data using the R package DESeq2. Then we conducted the MapMan enrichment analysis using MapMan4 (Schwacke et al., 2019), with the results of the gene differential expression analysis and the Mercator4 annotation as the input. For each gene, the functional annotation of the isoform with highest read count across all the samples was selected as the annotation of the gene. Graphs were plotted using the R package ggplot2.

**Measurements of phyllotaxis and data analysis.**

Plants for the phyllotaxis experiment were sampled at all three sites in the Biedouw valley described above, between 30 August and 9 September 2023.

Angles between subsequent leaf/floral stalk primordia were measured with a hinged protractor, a circular histogram that can be clipped non-destructively around the plant's stem. The circle was divided into 16 equal bins of 22.5°, with bin 1 centered around 0 degrees, and bin numbers increasing in a clockwise (CW) fashion. The angle of the leaf at the base of a floral stalk, when bent outwards, was used as proxy for primordium orientation. With the older organ aligned to the 0 degree line within bin 1, bin 11 corresponds to the 137.5° angle of a counterclockwise (CCW) phyllotactic spiral, when looking from older to younger organs, as is common in the field. Conversely, bin 7 corresponds to a clockwise golden angle. See Figure S2A. (Note: data was recorded young to old (bin 11 = CW), but transformed to old to young (bin 7 = CW) to avoid confusion.)

Histograms of subsequent flower angles were made from all data recorded before September 9. For this analysis, we only filtered against confounded developmental sequences (e.g., containing a bud in an older position than an open flower). All recorded angles are included in the analysis shown in Figure S2B (88 plants, 247 angles).

As in all but five sampled plants, the phyllotactic spiral was very consistent, i.e., either fully CW or fully CCW, we used a simplified scoring for the *C. alba* plants harvested on September 9: whole plants were scored as CW or CCW, without explicitly measuring the relevant angles. Individual flower orientations were scored as L (left), LU (left-unsure), U (unknown/impossible to determine), R (right) and RU (right-unsure). Where possible, buds were opened to determine flower orientation. If a plant displayed at least one L-flower and no R, the plant's flower orientation was scored as left (-1). In the opposite scenario, it was scored as right (1).

In all other cases, i.e., containing both orientations, or no L/R information, the plant was scored as mixed (0).

For phyllotactic orientation, the number of CCW (bins 2-8) and CW (bins 10-16) angles on a plant were counted. Bin 9 (angles of approximately 180°) was ignored. If both CW and CCW occurred on a single plant, its phyllotactic orientation was coded as unclear (0). We interpret these cases as analogous to the deviations from the regular phyllotactic spiral seen in *A. thaliana ahp6* mutants (so-called M-shapes) where the relative timing of organ initiation is changed, yet the spatial pattern maintained, resulting in three consecutive angles that are much larger than 137.5° (Besnard et al., 2014). Three of the five plants scored as having unclear phyllotaxis showed a deviation consistent with the beginning of an M-shape in their last measured divergence angle. As many plants had very short internodes (few mm or less), it was sometimes difficult to observe the correct developmental order of organs along the stem, so some of these observations could also have been caused by measurement errors. For similar reasons, plants on which the recorded sequence of generative organs contradicted normal developmental order (e.g., containing a bud in an older position than an open flower), were excluded from the filtered data set. Otherwise, all CW plants were coded 1 and all CCW plants were coded -1. In case relative angles of less than 90 degrees were recorded (bins 1-4 and 14-16), the phyllotactic pattern was considered confounded and these plants were not considered in the filtered data set. Filtering / preprocessing was done using a custom-built python script (processPhylloInfo.py).

Analysis of deviations from prediction relative to flower position was done using a custom built python script (findDeviationsFromPattern.py) and R (analyzePhyllotaxis.R). Correlation coefficients and 95% confidence intervals were calculated using the function corr.test in R. Based on the observed strong correlation between CW phyllotaxis and left-handed flowers, we predicted the flower orientation of all plants with consistent (fully CW or fully CCW) phyllotaxis. Per flower position, with 1 as the oldest primordium, the fraction of deviations (R on CW or L on CCW) was recorded. Confidence intervals for the fraction of deviations per position were calculated based on a binomial test. The baseline of uniform probability of deviations was computed as NumberOfFlowersThatDeviate / TotalNumberOfFlowersConsidered. Overall departure from this uniform prediction was tested using $X^2$ (chisq.test in R).

**Stereomicroscopy.**

We imaged dissected pistils from three different angles to obtain images of the three carpels that make up the ovary using a Nikon SMZ1500 Stereomicroscope fitted with a Nikon Digital Sight DS-Fi2 camera.

**Fluorescent microscopy.**

Fixation. We fixed whole pistils in 4% (w/v) paraformaldehyde (PFA) prepared in 1X phosphate buffered saline (PBS: 136.89 mM NaCl, 2.68 mM KCl, 5.37 mM $Na_2HPO_4$, 1.76 mM $KH_2PO_4$; pH7.4) for 1h at room temperature. Fixative was removed by rinsing three times in 1X PBS for 5 minutes, and once in $dH_2O$.

Clearing. We removed endogenous pigments using the ClearSee protocol for pistils described in Kurihara *et al.* (2015), with samples incubated in ClearSee (10% (w/v) Xylitol; 5% (w/v) sodium deoxycholate and 25% (w/v) Urea in $dH_2O$) for 6 weeks, or until transparent. The solution was changed three times per week until tissues had cleared.

Cell wall staining. Pistils were rinsed in $dH_2O$ for 1 h before being transferred into Calcofluor white solution (Sigma, product no. 18909) for overnight staining. Samples were de-stained overnight in $dH_2O$. All staining and de-staining steps were performed in the dark with gentle agitation. Samples were placed on long coverslips (25 X 50 mm) in water for imaging.

Imaging. Cell lengths of epidermal cells overlying the midveins of carpels 1 and 2

We performed fluorescent microscopy using a Zeiss LSM 880 Confocal and overview images of the entire pistil were taken using tile scanning on a widefield Zeiss monochrome CCD camera. We selected two positions on the ovary to take detailed z-stacks of cell structure. The first position was within the top third of the ovary, centred around one of the midveins that runs along the surface of each carpel. The second position was at the ovary-style transition, at the base of the style. The approximate positions of each z-stack were marked by eye on the overview image and the sample was then flipped using tweezers to image the opposite side, and the process repeated.

Imaging. Cell lengths of subepidermal cells in inner and outer walls of carpels 1 and 2

To obtain images of the inner walls of *C. alba* ovaries, we removed the adaxial surface of the ovary. Working under a dissecting microscope, we made longitudinal cuts along the prominent midveins of carpels one and two, as well as along either side of the indentation between the two adaxial carpels (i.e. along the carpel wall that separates carpel 1 from carpel 2). A final transverse cut along the base of the ovary was made and the two triangular pieces of tissue removed to reveal the chambers of the adaxial carpels. The ovules were removed by scraping along the placental wall with the tip of a scalpel blade.

Images of the dissected pistil were captured on a Zeiss LSM 880 Confocal microscope under 10X magnification, before taking higher resolution 20X Z-stack images of the cell walls of carpel 1 and carpel 2 at the same relative position.

### Image analysis and processing.

All image analysis was performed using the freely available online software ImageJ (https://ij.imjoy.io/).

Carpel lengths.

We measured carpel lengths using the spline tool. Each carpel was measured three times and the average recorded. Scale bars in each image were also measured in triplicate, using the straight-line measurement tool, and used to convert the length of the carpel in pixels to a measurement in micrometres. To account for variations in flower size, we normalized carpel lengths of individual samples against ovary size using the equation:

$$normalised\ length\ of\ carpel\ x = \frac{length\ of\ carpel\ x\ (\mu m)}{(\sum_1^3 length\ of\ carpel\ i)/3}$$

Cell lengths.

*Upper third of the ovary*: We chose an image from each z-stack that clearly showed the outlines of a line of cells running directly above each midvein. We then measured cell lengths of ten cells using the straight-line tool in ImageJ. We measured scale bars in each picture in triplicate and these were used to convert the mean cell length measurements in pixels to measurements in micrometres.

*Base of the style*: Z-stacks captured the ovary-style transition. For each carpel of the style, we chose a line of cells approximately 100 µm from the central crease of the style. An image which clearly captured a line of five or more cells in this position was exported and cell lengths were measured in ImageJ, as described above. We calculated averages over five cells rather than ten.

Cell widths.

Using the same images as for *Cell lengths: Upper third of the ovary*, we measured cell widths of five cells at 90° to the previous measurement. Cell measurements were performed as previously described.

Carpel outer and inner cell walls

An image from the z-stack with the clearest resolution was selected to quantify the lengths of 10 adjacent cells in the subepidermal layer of the outer and inner walls of carpels 1 and 2. These measurements were repeated three times for each image.

### Auxin treatment

Indole acetic acid (IAA; Sigma-Aldrich) was dissolved in absolute ethanol to produce a stock solution containing 100 mg/mL IAA. Lanolin containing auxin was prepared by melting 0.5 g of lanolin at 40°C and adding IAA stock solution to give a final concentration of either 5 mg/mL or 25 mg/mL IAA. After the IAA addition, the paste was mixed with a pipette tip, vortexed vigorously, and left to solidify. For the mock treatment, a corresponding volume of ethanol was mixed with lanolin.

*Cyanella alba flavescens* plants were collected from site 1 in the Biedouw Valley and maintained in water with cut flower food (Chrysal: Clear universal flower food). Individuals were selected to have at least one open flower and at least one early-stage bud for treatment. The handedness of the buds was extrapolated from the style deflection of the open flowers. We opened early-stage buds, removed two of the five adaxial stamens overlying carpel 2 and applied a small bead of lanolin paste to carpel 2 using a wooden toothpick. After closing the buds again the plants were kept at room temperature. When the treated flowers opened, all floral organs except the pistil were removed, the midvein of carpel 1 was marked and the pistils were photographed under a dissecting microscope (Olympus SZ61) fitted with a Zeiss Axiocam 208 colour camera from the adaxial side, facing the line where carpels 1 and 2 are fused (see Stereomicroscopy section above). From these images we measured the lengths of midveins 1 and 2 as above. Style deflection was quantified by measuring the angle between a line drawn perpendicular to the base of the ovary and a line drawn through the middle of the style at its base.

**Supporting Information References:**

Besnard, F., Refahi, Y., Morin, V., Marteaux, B., Brunoud, G., Chambrier, P., Rozier, F., Mirabet, V., Legrand, J., Lainé, S., Thévenon, E., Farcot, E., Cellier, C., Das, P., Bishopp, A., Dumas, R., Parcy, F., Helariutta, Y., Boudaoud, A., Godin, C., Traas, J., Guédon, Y., Vernoux, T., 2014. Cytokinin signalling inhibitory fields provide robustness to phyllotaxis. Nature 505, 417–421. https://doi.org/10.1038/nature12791

Cheng, H., Concepcion, G.T., Feng, X., Zhang, H., Li, H., 2021. Haplotype-resolved de novo assembly using phased assembly graphs with hifiasm. Nat. Methods 18, 170–175. https://doi.org/10.1038/s41592-020-01056-5

Danecek, P., Auton, A., Abecasis, G., Albers, C.A., Banks, E., DePristo, M.A., Handsaker, R.E., Lunter, G., Marth, G.T., Sherry, S.T., McVean, G., Durbin, R., 1000 Genomes Project Anal Grp, 2011. The variant call format and VCFtools. Bioinformatics 27, 2156–2158. https://doi.org/10.1093/bioinformatics/btr330

Danecek, P., Bonfield, J.K., Liddle, J., Marshall, J., Ohan, V., Pollard, M.O., Whitwham, A., Keane, T., McCarthy, S.A., Davies, R.M., Li, H., 2021. Twelve years of SAMtools and BCFtools. GigaScience 10, giab008. https://doi.org/10.1093/gigascience/giab008

Dobin, A., Davis, C.A., Schlesinger, F., Drenkow, J., Zaleski, C., Jha, S., Batut, P., Chaisson, M., Gingeras, T.R., 2013. STAR: ultrafast universal RNA-seq aligner. Bioinformatics 29, 15–21. https://doi.org/10.1093/bioinformatics/bts635

Gabriel, L., Brůna, T., Hoff, K.J., Ebel, M., Lomsadze, A., Borodovsky, M., Stanke, M., 2023. BRAKER3: Fully automated genome annotation using RNA-Seq and protein evidence with GeneMark-ETP, AUGUSTUS and TSEBRA. BioRxiv Prepr. Serv. Biol. 2023.06.10.544449. https://doi.org/10.1101/2023.06.10.544449

Grabherr, M.G., Haas, B.J., Yassour, M., Levin, J.Z., Thompson, D.A., Amit, I., Adiconis, X., Fan, L., Raychowdhury, R., Zeng, Q., Chen, Z., Mauceli, E., Hacohen, N., Gnirke, A., Rhind, N., di Palma, F., Birren, B.W., Nusbaum, C., Lindblad-Toh, K., Friedman, N., Regev, A., 2011. Full-length transcriptome assembly from RNA-Seq data without a reference genome. Nat Biotechnol 29, 644–52. https://doi.org/10.1038/nbt.1883 nbt.1883 [pii]

Kokot, M., Dlugosz, M., Deorowicz, S., 2017. KMC 3: counting and manipulating k-mer statistics. Bioinforma. Oxf. Engl. 33, 2759–2761. https://doi.org/10.1093/bioinformatics/btx304

Kuznetsov, D., Tegenfeldt, F., Manni, M., Seppey, M., Berkeley, M., Kriventseva, E.V., Zdobnov, E.M., 2023. OrthoDB v11: annotation of orthologs in the widest sampling of organismal diversity. Nucleic Acids Res. 51, D445–D451. https://doi.org/10.1093/nar/gkac998

Love, M.I., Huber, W., Anders, S., 2014. Moderated estimation of fold change and dispersion for RNA-seq data with DESeq2. Genome Biol 15, 550. https://doi.org/s13059-014-0550-8 [pii] 10.1186/s13059-014-0550-8

Manni, M., Berkeley, M.R., Seppey, M., Simão, F.A., Zdobnov, E.M., 2021. BUSCO Update: Novel and streamlined workflows along with broader and deeper phylogenetic coverage for scoring of eukaryotic, prokaryotic, and viral genomes. Mol. Biol. Evol. 38, 4647–4654. https://doi.org/10.1093/molbev/msab199

Manning, J.C., Goldblatt, P., 2012. A revision of Tecophilaeaceae subfam. Tecophilaeoideae in Africa. Bothalia Afr. Biodivers. Conserv. 42.

Mikheenko, A., Prjibelski, A., Saveliev, V., Antipov, D., Gurevich, A., 2018. Versatile genome assembly evaluation with QUAST-LG. Bioinforma. Oxf. Engl. 34, i142–i150. https://doi.org/10.1093/bioinformatics/bty266

Patro, R., Duggal, G., Love, M.I., Irizarry, R.A., Kingsford, C., 2017. Salmon provides fast and bias-aware quantification of transcript expression. Nat. Methods 14, 417–419. https://doi.org/10.1038/nmeth.4197

Pedersen, B.S., Quinlan, A.R., 2018. Mosdepth: quick coverage calculation for genomes and exomes. Bioinforma. Oxf. Engl. 34, 867–868. https://doi.org/10.1093/bioinformatics/btx699

Pertea, M., Pertea, G.M., Antonescu, C.M., Chang, T.-C., Mendell, J.T., Salzberg, S.L., 2015. StringTie enables improved reconstruction of a transcriptome from RNA-seq reads. Nat. Biotechnol. 33, 290–295. https://doi.org/10.1038/nbt.3122

Quinlan, A.R., Hall, I.M., 2010. BEDTools: a flexible suite of utilities for comparing genomic features. Bioinformatics 26, 841–2. https://doi.org/10.1093/bioinformatics/btq033 btq033 [pii]

Ranallo-Benavidez, T.R., Jaron, K.S., Schatz, M.C., 2020. GenomeScope 2.0 and Smudgeplot for reference-free profiling of polyploid genomes. Nat. Commun. 11, 1432. https://doi.org/10.1038/s41467-020-14998-3

Schwacke, R., Ponce-Soto, G.Y., Krause, K., Bolger, A.M., Arsova, B., Hallab, A., Gruden, K., Stitt, M., Bolger, M.E., Usadel, B., 2019. MapMan4: A refined protein classification and annotation framework applicable to multi-omics data analysis. Mol. Plant, Plant Systems Biology 12, 879–892. https://doi.org/10.1016/j.molp.2019.01.003

Sedlazeck, F.J., Rescheneder, P., von Haeseler, A., 2013. NextGenMap: fast and accurate read mapping in highly polymorphic genomes. Bioinforma. Oxf. Engl. 29, 2790–2791. https://doi.org/10.1093/bioinformatics/btt468

Thorvaldsdottir, H., Robinson, J.T., Mesirov, J.P., 2013. Integrative Genomics Viewer (IGV): high-performance genomics data visualization and exploration. Brief. Bioinform. 14, 178–192. https://doi.org/10.1093/bib/bbs017

Wang, Y., Chen, Q., Deng, C., Zheng, Y., Sun, F., 2020. KmerGO: A tool to identify group-specific sequences with k-mers. Front. Microbiol. 11, 2067. https://doi.org/10.3389/fmicb.2020.02067

Wickham, H., 2009. ggplot2: Elegant Graphics for Data Analysis. Springer-Verlag, New York.
